# Predominantly genetic determination and stable transmission of DNA methylation in an avian hybrid zone

**DOI:** 10.64898/2026.02.27.708517

**Authors:** Fritjof Lammers, Valentina Peona, Madeline Chase, Dave Lutgen, Marta Burri, Reto Burri

## Abstract

The reshuffling of divergent genomes upon hybridization may disrupt co-evolved regulatory systems and contribute to epigenetic instability and, ultimately, reproductive isolation. While the genetic consequences of hybridization are well documented, insights into the consequences of hybridization for DNA methylation are currently limited. To obtain insights into the regulation of methylation and its transmission under hybridization, we here investigated genome-wide methylation in a natural hybrid zone of songbirds (wheatears of the *Oenanthe hispanica* complex) by integrating nearly 100 methylomes with population genomic data. Across 436,762 CpG sites, the population structure of methylation closely mirrors genetic population structure. Methylation quantitative trait locus analyses identify widespread associations of genetic with methylation variation, predominantly in trans, consistent with a regulatory architecture in which the genetic background determines methylation variation. Between species, methylation divergence is limited, with only 0.31% of CpGs differentially methylated. While at the level of chromosomes methylation divergence strongly correlates with genetic differentiation, the extent to which differentially methylated loci coincide with high genetic differentiation differs among chromosomes. A close-to-absent methylation divergence from promoters and coding regions indicates conservation of core regulatory architectures. Finally, CpGs with highest methylation divergence exhibit predominantly additive or dominant transition patterns in hybrids. In contrast, transgressive methylation is exceedingly rare, and we find no evidence for widespread hybrid-induced demethylation. Or results corroborate that DNA methylation primarily reflects underlying genetic variation in birds and remains robust to genome reshuffling, and at least for wheatears suggest a limited role for methylation divergence in hybrid dysfunction and reproductive isolation.

## Introduction

Hybridization is a widespread evolutionary process that shapes biodiversity by bringing together genomes that previously evolved in isolation (Abbott et al. 2013; Taylor and Larson 2019). By combining divergent genomes, hybridization reshuffles genetic variation and regulatory architectures into new combinations. Thereby, this genomic admixture can generate novel phenotypic diversity (Abbott et al. 2013) and by exposing genetic incompatibilities between genomic regions contribute to reproductive isolation (Presgraves 2010; Moran et al. 2024). Therefore, hybridization provides a powerful framework to study how regulatory systems respond to the admixture of divergent genomes (Wolf et al. 2010).

While the genetic consequences of hybridization have been studied extensively (Moran et al. 2021), its effects on gene regulatory systems remain less well understood (Runemark et al. 2024). Studies in Drosophila (Wittkopp et al. 2004) and mice (Goncalves et al. 2012) show that the divergence in cis-regulatory elements and trans-acting factors can lead to altered gene expression and regulatory incompatibilities in hybrids (Michalak 2009; Mack and Nachman 2017; Runemark et al. 2024). Yet it remains unclear whether epigenetic regulatory layers that shape gene expression are similarly disrupted by genomic admixture (Berbel-Filho et al. 2022) or whether they remain constrained by genome-wide ancestry and are stably transmitted, similar to the inheritance of genetic diversity (Merondun and Wolf 2025). Among epigenetic regulatory layers, DNA methylation (hereafter ‘methylation’) represents a widely conserved and pervasive epigenetic modification that has strong regulatory potential (Bird 2002), important roles in maintaining genome stability (Walter et al. 2016), and possibly important role in reproductive isolation and speciation (Smith et al. 2016; Laporte et al. 2019).

In vertebrates, methylation primarily occurs as 5-methylcytosine (5mC) in CpG context and plays a central role in transcriptional regulation, development, and genome stability (Bird 2002; Chapelle and Silvestre 2022). Although methylation patterns can respond to developmental and environmental variation (Sheldon et al. 2020), population-scale studies indicate that much of the individual variation in methylation is shaped by underlying genetic variation (McRae et al. 2014; Wang et al. 2016; Merondun and Wolf 2025) that usually persists even under developmental and environmental disturbance (Sepers et al. 2023a, 2023b).

Despite increasing evidence for genetic determination of methylation, it remains unclear how robust the genetic determination of extensively methylation is when divergent genomes recombine upon hybridization. In marsupials and *Coregonus* fish, hybridization induces hypomethylation of transposable elements (O’Neill et al. 1998; Laporte et al. 2019), possibly caused by the disruption of co-adapted regulators that are exposed to novel genomic combinations. Alternatively, methylation patterns may remain tightly constrained by the genome-wide genetic background even under extensive admixture (Berbel-Filho et al. 2022; Merondun and Wolf 2025). Two expectations can therefore be formulated for transmission patterns of methylation in hybrid genomes. Under a model of primarily constrained genetic determination, methylation variation is strongly correlated to genome-wide genetic ancestry, leading to additive or dominant methylation patterns in hybrids and limited hybrid-specific methylation patterns that would hint towards transgressive effects of new genomic combinations on methylation regulation (Kakoulidou and Johannes 2024). Alternatively, if the regulation of methylation is disrupted by recombination of formerly co-inherited haplotypes, hybrid-specific, respectively transgressive methylation signatures may emerge (O’Neill et al. 1998; Laporte et al. 2019).

The *Oenanthe hispanica* species complex (hereafter ‘*hispanica* complex’) of wheatears provides an ideal system to test these predictions in a natural setting. Within this songbird complex, the Western Black-eared Wheatear (*Oenanthe melanoleuca*) and the Pied Wheatear (*O. pleschanka*) form a stable hybrid zone in northern Iran that is characterized by extensive admixture and gradual transitions in genetic ancestry (Schweizer et al. 2019; Lutgen et al. 2025). The presence of the entire spectrum of admixture provides a powerful system to study the transmission of methylation upon hybridization. Here, we sequenced population-scale whole-methylomes of >100 individuals across this admixture spectrum and combined them with matching whole-genome sequencing data to investigate how methylation varies between parental species and across hybrids in this natural hybrid zone. By integrating methylation profiles with measures of genetic ancestry and genomic differentiation, we test how methylation is transmitted in a hybridizing species and whether hybridization induces a disruption of methylation regulation.

Specifically, we address three questions: 1. Does methylation variation in the *hispanica* complex and the hybrids between *O. melanoleuca* and *O. pleschanka*, reflect genetic ancestry, that is, is methylation variation determined predominantly by genetic variation? 2. How divergent are methylomes between *O. melanoleuca* and *O. pleschanka*, and how does methylation divergence relate to genetic differentiation? 3. How does hybridization affect methylation states: do hybrids predominantly exhibit additive or dominant transmission patterns consistent with individuals’ genetic ancestry, or do they show evidence of transgressive methylation? Our results demonstrate a predominant contribution of genetic regulation to methylation variation within the *hispanica* complex, reflected in highly congruent population structure of methylation and genetic variation between populations. The two parental species exhibit partially co-localized signatures of genetic and methylation differentiation. Hybrids exhibit mostly additive or dominant methylation signatures that reflect hybrids’ ancestry compositions. Transgressive methylation patterns are exceedingly rare in wheatear hybrids, suggesting that hybridization does not broadly disrupt methylation regulation in this avian system.

## Results

### Data description

To analyze methylation profiles in O. *melanoleuca*, O. *pleschanka*, and their hybrids from the Iranian hybrid zone, we generated population-scale whole-genome methylation data for *O. melanoleuca* (n=19), *O. pleschanka* (n=12), their hybrids (n=56), and for related species in the *O. hispanica* complex (**fig. 1A,B**; **tables S1, S2**). After creating individualized reference genomes using whole-genome sequencing data of the same samples to account for C>T (and G>A) polymorphisms (see Materials and Methods), quality-trimmed sequencing reads mapped with a mean coverage of 12.82 ± 1.9× and an alignment rate of 70.9 ± 3.9%. From these alignments, Bismark called methylation levels from 6.98 ± 0.19M CpG sites per sample, out of which we retained 436,762 CpG sites with >5× coverage in 98 samples and no missing values for further analysis. Three samples were discarded due to insufficient sequence data quality (**table S2**).

**Figure 1.**
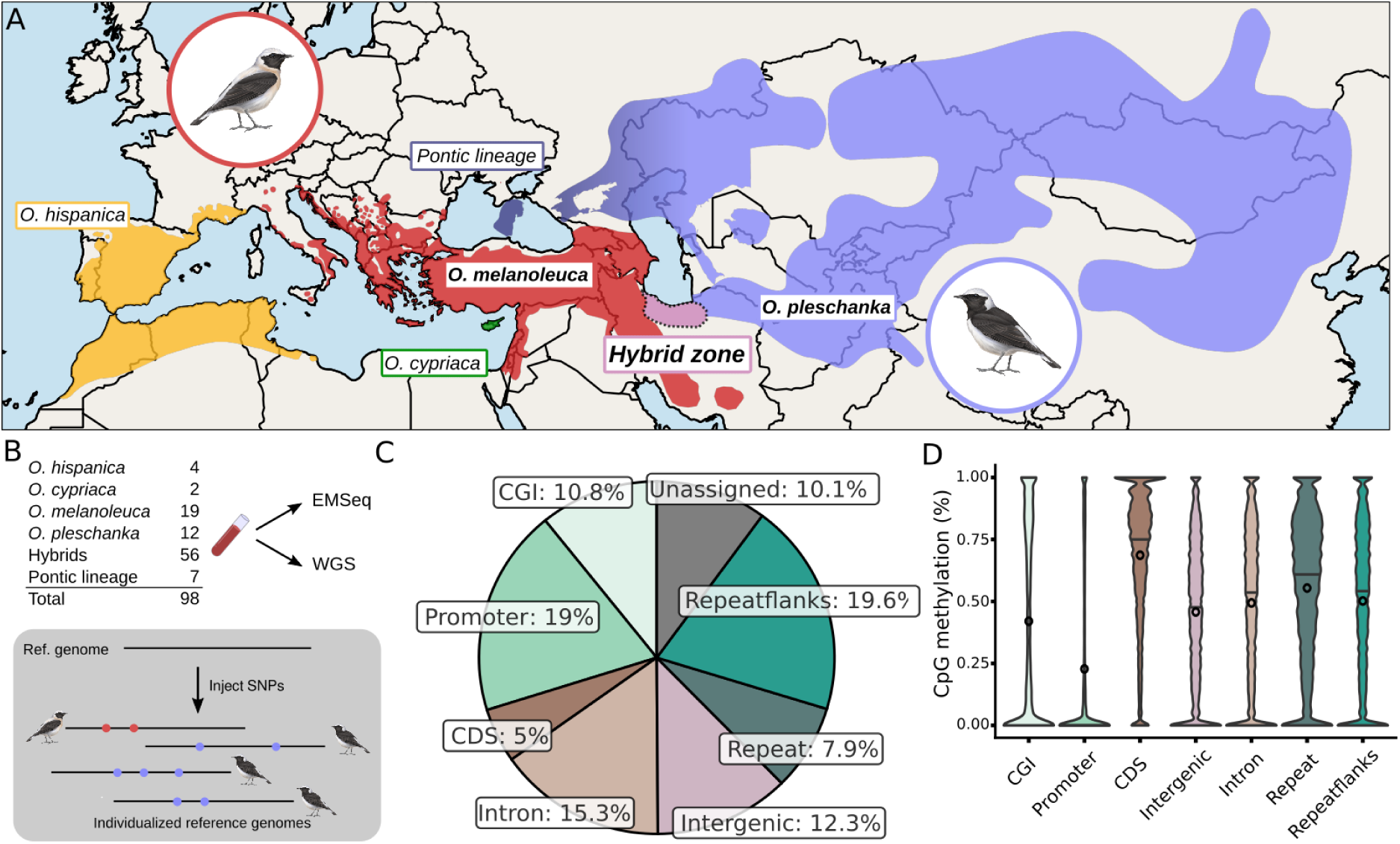
Sampling design and dataset structure. **A)** Geographic distribution of sampled populations in the *Oenanthe hispanica* complex. Primarily, samples from *O. melanoleuca*, *O. pleschanka* and the hybrid zone occurring in northern Iran were sequenced. **B)** Methods summary with sample sizes per population. Methylomes were sequences from blood using EMSeq and complemented with available whole-genome sequencing (WGS) from a previous study. To improve genome alignments, and account for C>T(G>A) mutations, individualized reference genomes were created using SNP calls from WGS data. **C)** Distribution of CpG sites in the *O. melanoleuca* reference genome across genomic contexts. **D)** Mean methylation levels across genomic contexts. Abbreviations: CGI, CpG islands; CDS, Coding Sequences; IG, Intergenic; Flanks, Repeat flanks.

Methylation patterns derived from these data were consistent both across the data set and with methylation patterns known from other vertebrates. First, within the data set, genome-wide methylation was consistent across samples and populations. Across all samples, the mean genome-wide methylation level was 0.521±0.045, with minor differences between populations (**supplementary table S2, supplementary fig. S1**). Based on our annotation of all 7.9M CpG sites in the OenMel1.1 reference genome, 19.6% of CpG sites were found in flanks of repetitive elements, followed by promotor-specific CpG islands (CGI; 19.0%), and introns (15.3%) (**fig. 1C**). Second, methylation levels varied strongly between genomic contexts, with the lowest methylation found in promoter-specific CpG islands (CGI, mean: 21.6%, median: 0.00%) and highest in coding sequences (mean: 68.7%, median: 75.6%; **fig. 1D**). These patterns of methylation levels across the genome match methylation patterns reported from other vertebrate genomes (Al Adhami et al. 2022).

In summary, our dataset reiterates properties of methylation biology established in vertebrate genomes, giving us confidence in our data and processing pipeline.

### Genetic background predicts methylation structure

First, to characterize genome-wide patterns of methylation variation across the investigated species and their hybrids, we applied a principal component analysis (PCA) on 436,762 CpG sites that were fully covered in all samples (no missingness). The first principal component (PC1; 10.2% variance explained) was correlated to mean genome-wide methylation level (R^2^=0.484, p=5.97^-15^) and clustered samples irrespectively to other biological or technical variation (**supplementary fig. 2**), indicating that methylation variance is similar across samples and constitutes a large fraction of variation in the dataset. Subsequent principal components (PCs) captured variation that reflected the geographic origin and genetic ancestry of individuals. We found that 8.6% of all variance tracked the geographic origin of individuals of *O. melanoleuca*, O. *pleschanka* and their hybrids by placing the two parental lineages from the opposite ends of the east-west gradient at the extremes of PC2 (**fig. 2A)**. In-between, the Iranian hybrids formed a gradient tracing their sampling location across northern Iran (see below). PC3 and PC4 captured additional variance (5.4% and 5.3%) that separated individuals from the Pontic lineage and *O. hispanica* from the *O. melanoleuca-O. pleschanka* cline (**supplementary fig. S2**). The Pontic lineage is a sister lineage to *pleschanka* from the main species range that has distinct demographic history and occurs in the Pontic steppes (Lutgen et al. 2025).

**Figure 2.**
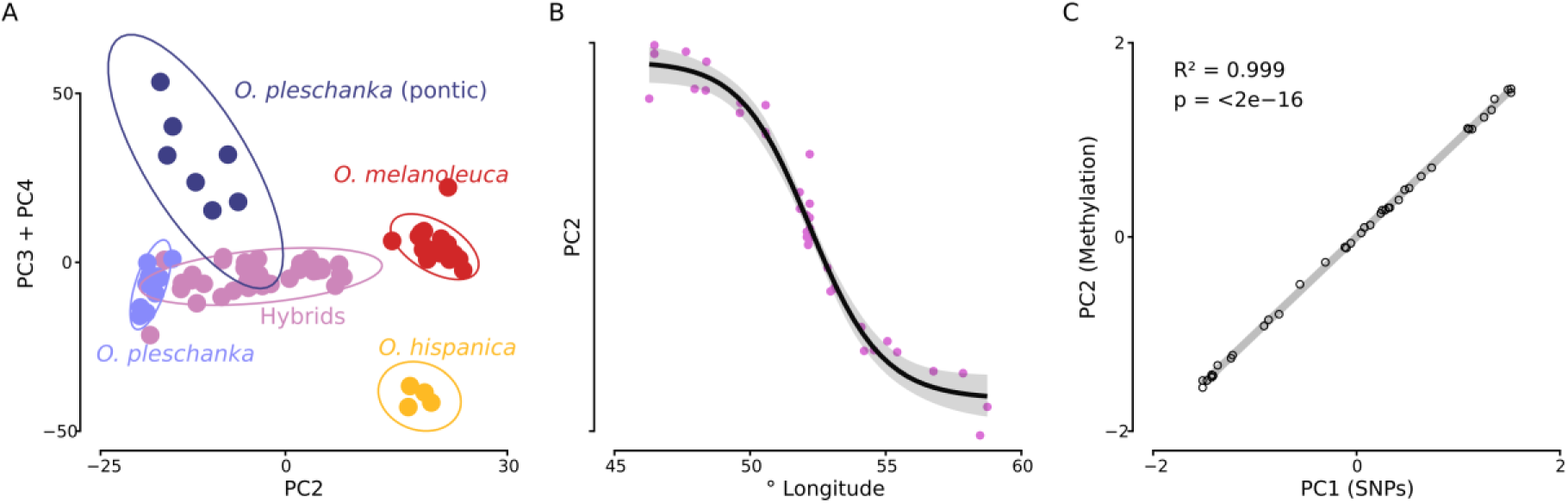
Structure of methylation variation the *hispanica* complex separates populations and is tightly linked to geographic origin of samples and their genetic structure. **A)** PCA of methylation data colored by genetically assigned population. **B)** Geographic structure of methylation across the hybrid zone. Shown is methylation PC2 against sample longitude with regression line and standard error shaded in grey (pseudo-R² = 0.97; residual SE = 1.39). **C)** Correlation of geographic structure of methylation and genetic variation. Methylation PC2 and genetic PC1 show a near perfect correlation. Both axes are Z-transformed for better comparability.

Because methylation patterns showed a strong association with individuals’ geographic origin and genetic ancestry, we next characterized the relationship of PCs with these variables. Across the Iranian hybrid zone PC2 strongly correlates with longitude, following a sigmoidal pattern (four-parameter logistic model; pseudo-R² = 0.97; residual SE = 1.39; **fig. 2B**) similar to the one previously described for genetic variation (Schweizer et al. 2019; Lutgen et al. 2025). To test whether genome-wide methylation variation correlates with genetic variation, we modeled methylation PC2 as a function of genome-wide genetic ancestry of the same samples (Lutgen et al. 2025). We found a near-perfect correlation between methylation and genetic ancestry (R^2^=0.99, p<2^-16^, **fig. 2C**). This suggests a strong genetic contribution of genetic variation to the determination of methylation variation in wheatears.

Given this indirect evidence for the genetic determination of methylation in *O. melanoleuca*, *O. pleschanka* and their hybrids, we next aimed to identify genotypes that underlie methylation states. To this end, we performed a methylation quantitative trait locus (meQTL) analysis. The meQTL analyses provide a genome-wide characterization of genotype-methylation associations within the constraints of the available sample size and population structure (Mueller et al. 2025). While these factors limit statistical power and result in inflated p-value distributions (**supplementary fig. S3**) and may underestimate the number of significant associations, they nonetheless enable an informative assessment at the level of summary statistics. We therefore restricted downstream interpretation to patterns that are robust to these limitations, rather than focusing on individual meQTLs. For the selection of genetic variants, we employed two different selection criteria. The first analysis, involving all available genetic variants (n=7,196,229) across *O. melanoleuca*, *O. pleschanka* and hybrids yielded 150,683 meQTLs, of which 17,674 (11.7%) were situated up to 1 million base pairs (Mb) away from CpGs and thus considered *cis*-associated and 133,009 (88.3%) were found in *trans* (>1 Mb). These 150,683 meQTLs were associated with 12,939 CpG sites, indicating this analysis was only able to associate a small subset of all CpGs (3.4% of variable CpG sites) with genetic variation. In addition, methylation levels at these CpG sites were correlated with genotypes at multiple SNPs (multi-SNP meQTLs, n=144,717; mean=20.8+-58.7 SNPs/CpG; **supplementary table S3**). We found similar results when repeating the meQTL analysis using an LD-pruned set of SNPs (n=3,797,382) that identified a third of the initially identified meQTLs (n=48,672, 77.0% overlapping) but shared 89.1% of CpG sites identified in the previous analysis. This suggests that linked SNPs contribute equally to methylation at associated CpG sites, that is, linked SNPs may act as a single meQTL.

Collectively, the genome-wide correspondence between genetic and epigenetic variation, as measured using methylation, and the predominance of *trans*-associated meQTLs and multi-SNP meQTLs indicate that genome-wide genetic ancestry, in addition to local genetic variation, strongly determines methylation states in the *hispanica* complex.

### Quantification of methylome divergence between species

To establish baseline patterns of methylation divergence between *O. melanoleuca* and *O. pleschanka* and provide a reference point for subsequent hybrid-focused analyses, we next aimed to characterize methylation divergence between the two parental species. Using beta-binomial regressions implemented in DSS (Feng and Wu 2019), we identified 1,396 differentially methylated loci (DMLs) and 37 differentially methylated regions (DMRs; comprising 283 CpGs) between the two species after applying a minimum methylation difference threshold of 20% and correcting for multiple testing (FDR < 0.01; **fig. 3A, supplementary fig. S4**). DMLs and DMRs represent 0.31%, respectively 0.064% of the 436,762 screened CpG sites. This suggests overall low to moderate levels of methylation divergence. Because DMLs were enriched on chromosome Z, which is present in a single copy in females, we excluded two female *O. pleschanka* samples and report results from the male-only analyses.

**Figure 3.**
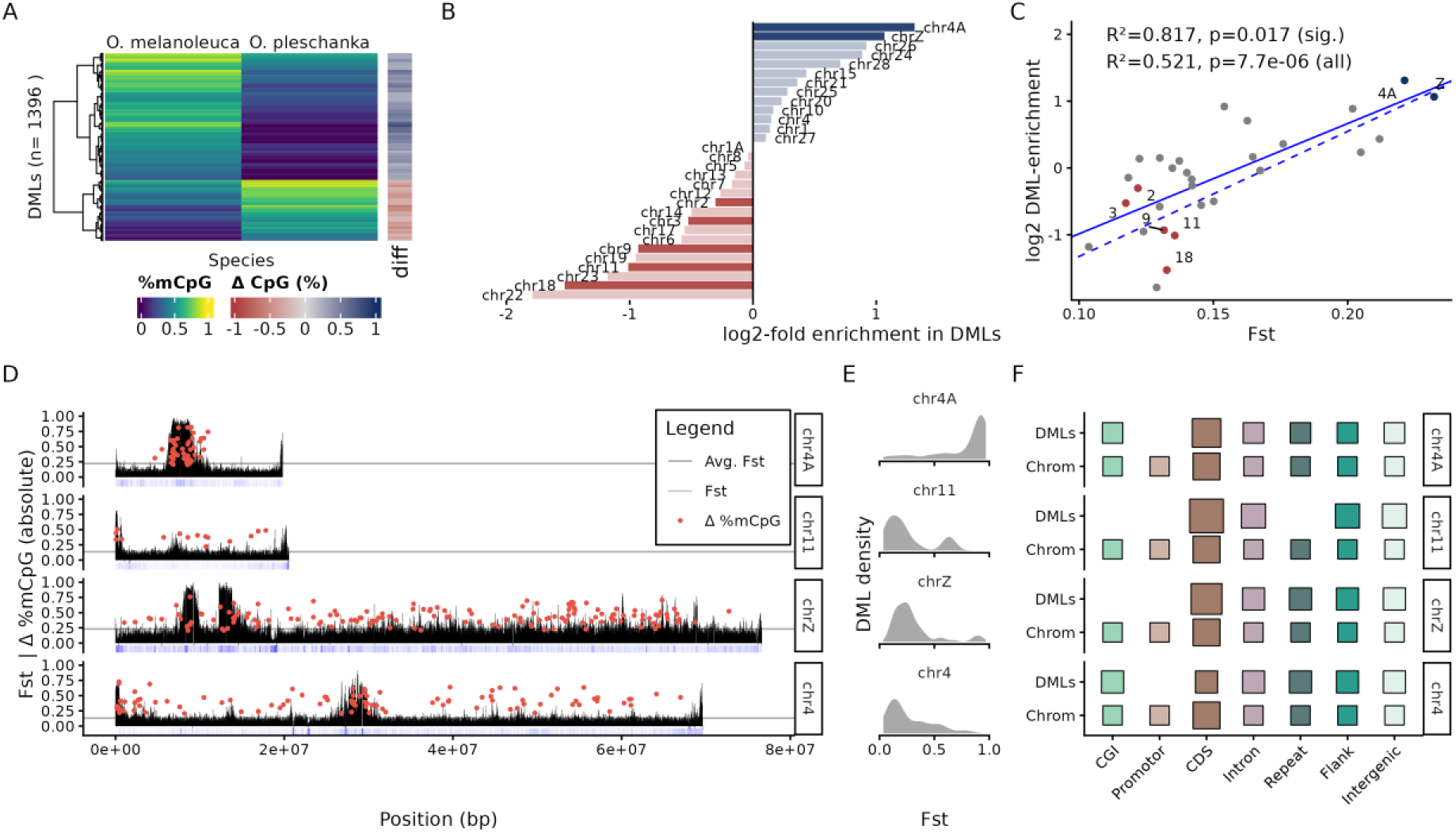
Methylation divergence between the parental species O. *melanoleuca* and O. *pleschanka* is non-randomly distributed and correlates with genetic differentiation (*F*_ST_). **A)** Heatmap of differentially methylated loci (DMLs) showing methylation level differences between *O. melanoleuca* and *O. pleschanka*. **B)** Enrichment and depletion of DMLs across chromosomes shows significant enrichment for chromosomes 4A and Z. Non-significant changes are shown in opaque colors. **C)** Chromosomal DML enrichment correlates with genetic differentiation (*F*_ST_). Linear regressions were calculated using all chromosomes (all, solid line) and chromosomes with significant changes only (sig., dashed line). **D)** Genetic differentiation and DMLs along highly DML-enriched chromosomes 4A and Z, as well as chromosomes 11 and 4 as comparisons. *F*_ST_ is shown in 10kb windows and locations of DMLs are indicated by red dots. **E)** DML distribution across the genetic differentiation (*F*_ST_) spectrum. DMLs coincide markedly with high-*F*_ST_ regions on chromosome 4A, whereas no such co-localization is present in other chromosomes, including chromosome Z. **F)** Comparison of DML occurrence (DMLs) with chromosome-wide CpG abundance (Chrom.) per genomic context for the four example chromosomes. Promoters, and partially also non-promotor CpG islands (CGI) are virtually absent from DMLs.

To assess how methylation divergence is distributed along the genome, we tested whether there were chromosomes enriched for DMLs. Based on the 436,762 DMLs included in the screen, we found a non-random distribution of DMLs across chromosomes (χ² = 241.44, df = 29, p < 2.2 × 10⁻¹⁶), with several chromosomes showing significant enrichment or depletion of DMLs (**fig. 3B**). Chromosomes 4A and Z showed the strongest enrichment (4A: log₂[obs/exp] = 1.31, FDR = 5.5 × 10⁻¹¹; Z: log₂[obs/exp] = 1.07, FDR = 2.32 × 10⁻¹⁷), and additional ten chromosomes exhibited non-significant trends toward an enrichment of DMLs (**fig. 3B**). Seventeen chromosomes were depleted of DMLs, with chromosomes 2, 3, 9, 11 and 18 showing significant depletion. A PCA excluding chromosomes Z and 4A asserted that methylation structure as reported above was not driven by the highly differentiated chromosomes (**supplementary fig. S5**).

Notably, chromosomes Z and 4A are also the two most genetically differentiated chromosomes between *O. melanoleuca* and *O. pleschanka*, as measured by the fixation index (*F*_ST_) (**fig. 3C**). Consistent with this observation, chromosome-wide average *F*_ST_ is a strong predictor of significant DML enrichment (R² = 0.817, p = 0.017; **fig. 3C**). This suggests that methylation divergence largely mirrors genetic differentiation at the chromosomal level. When chromosomes without significant DML enrichment were included in the statistical model, the relationship was weaker but remained significant (R² = 0.521, p = 7.7 · 10^-6^; **fig. 3C**), indicating a genome-wide correlation of methylation divergence with genetic differentiation.

Observing that chromosomes differ markedly in their overall degree of methylation divergence, we next examined the distribution of DMLs and *F*_ST_ along chromosomes. Mapping *F*_ST_ in 10kb windows together with the coordinates of DMLs revealed that chromosomes show distinctive patterns of co-localization of methylation divergence and genetic differentiation. This co-localization was highly distinct between the two chromosomes showing the largest overall levels of methylation differentiation, chromosomes 4A and Z (**fig. 3D, supplementary fig. S6**). On chromosome Z, DMLs are broadly distributed along the chromosome and show no apparent co-localization with genetic differentiation, whereas chromosome 4A exhibits a pronounced peak where elevated genetic differentiation and DMLs coincide (**fig. 3D, E)**. The association of DML density with high-*F*_ST_ regions on chromosome 4A is also apparent in density plots and starkly contrasts the patterns observed on other chromosomes, including chromosome Z, which are highly depleted of DMLs in regions of medium (*F*_ST_>0.3) to high (*F*_ST>_ 0.7) genetic differentiation (**fig. 3E**, **supplementary fig. S7**). This difference is not explained by chromosome length, as comparisons with similar-sized chromosomes 11 and 4 show (**fig. 3D**). Instead, we find that chromosome 4A contains no intrachromosomal trans-associated meQTLs, and the ratio of cis and trans-associated meQTLs was significantly different between chromosome 4A and Z (χ² = 23.443, df = 1, p-value = 1.286e-06), even after correcting for different chromosome length (χ² = 7.4453, df = 1, p-value = 0.0063). Thus, the contrasting patterns of chromosome-wide methylation divergence and, in particular, its co-localization with genetically differentiated regions on chromosome 4A, suggest that regulatory architectures of methylation can differ across the genome.

Finally, we assessed whether methylation divergence between our study species is subject to different levels of constraint, depending on the affected genomic region’s function. To this end, we tested whether inter-species DMLs were enriched in regulatory or coding regions, in which methylation is most likely to have effects on the phenotype. When comparing the occurrence of DMLs across genomic contexts, we found no enrichment of DMLs in promoters, coding sequences, or other regulatory contexts (**fig. 3F**; **supplementary fig. S8**). Instead, across most chromosomes, DMLs were virtually absent from promoters and within coding sequences showed only minor deviations from expectations. These results suggest that core gene regulatory architecture mediated by methylation is largely conserved between *O. melanoleuca* and *O. pleschanka*.

In summary, our analyses reveal generally low to moderate methylation divergence between *O. melanoleuca* and *O. pleschanka*, with pronounced heterogeneity across chromosomes. Elevated methylation divergence on chromosomes Z and 4A coincides with strong genetic differentiation even when accounting for potentially confounding properties, such as GC content or repeat abundance (**fig. 3C**). Yet these chromosomes differ markedly in the spatial correlation of genetic differentiation and methylation divergence, implying different regulatory architectures or evolutionary processes underlying methylation divergence within the same genome. The absence of methylation divergence in regulatory regions further suggests that gene expression programs are broadly conserved between the two species investigated here, and that methylation divergence predominantly reflects underlying genomic differentiation rather than independent evolution of methylation.

### Stable transmission of methylation in hybrids, and minimal transgressive methylation patterns

Hybrids provide a natural experiment that enables us to assess how methylation patterns evolve as they expose divergent methylation variation to novel genetic backgrounds and environments (Mueller et al. 2025). Following this idea, we leveraged methylation variation in the hybrids between O. *melanoleuca* and O. *pleschanka* to understand how the reshuffling of genetic ancestries may affect the regulation of methylation, and, notably, test whether hybridization induces transgressive methylation that may contribute to novel hybrid phenotypes (Lauss et al. 2018).

First, we characterized how hybrid methylomes differ from their parental populations by comparing their genome-wide methylation levels and performing screens for DMLs between hybrids and the parental species. In hybrids, the average genome-wide methylation was slightly reduced (50.3±1.75% vs 50.7±2.33% and 50.8±2.21%) but not significantly different from parental species (**fig. 4A**, one-sided Wilcoxon rank sum exact test, p=0.099). To find potential site-specific methylation differences between hybrids and each parental species, we inferred DMLs. In addition to the previously identified 1,396 DMLs between parental species (inter-species DMLs’), we detected 375 DMLs between *O. melanoleuca* and hybrids (247 shared with inter-species-DMLs, 76 unique, 54 shared across all comparisons) and 215 DMLs between O. *pleschanka* and hybrids (131 shared with inter-species-DMLs, 30 unique, 54 shared across all comparisons). Of the 1,396 inter-species-DMLs, 441 overlapped with at least one hybrid comparison (**fig. 4B**). The largest overlap was observed between inter-species and *melanoleuca*-hybrid DMLs (n = 247, 16%), indicating stronger methylation divergence between *O. melanoleuca* and hybrids than between *O. pleschanka* and hybrids (n = 131, 9%). Only a small fraction of DMLs was unique to comparisons involving hybrids and either parental species (5% and 2%, respectively), suggesting that methylation divergence specific to hybrids (that is, methylation diverged in hybrids from both parental species) is very limited.

**Figure 4.**
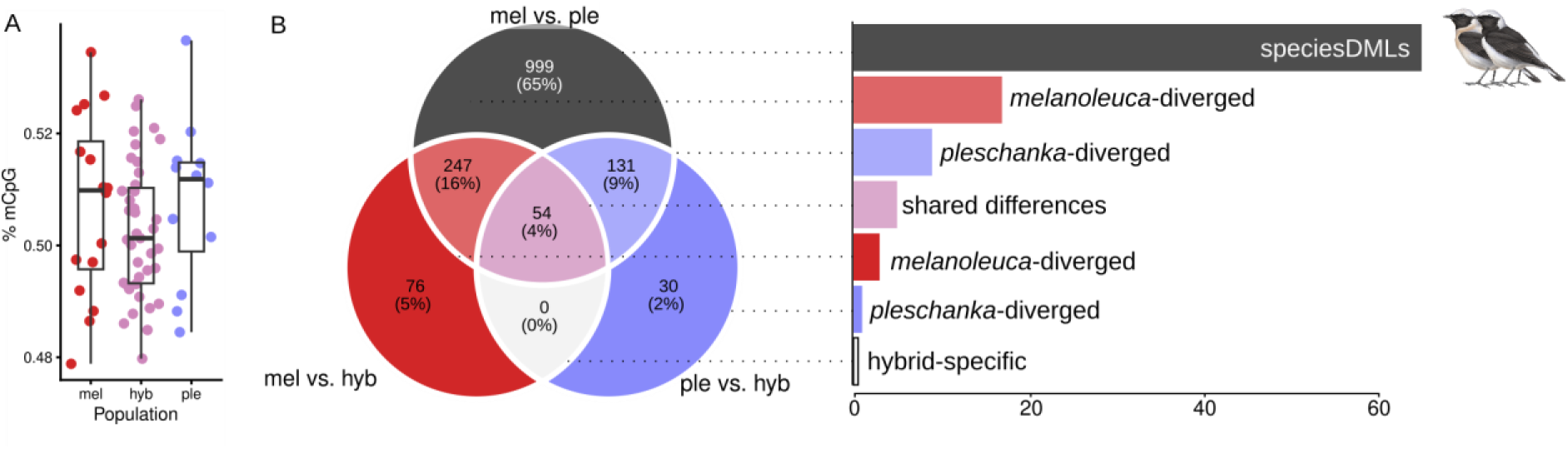
Methylation divergence in hybrids. **A)** Genome-wide mean methylation levels show a non-significant trend to greater variance in hybrids compared to the parental species (permutation-test 1,000 replicates, test-statistic=-0.001, p=0.79). Population differences are minimal and non-significant (one-sided Wilcoxon rank sum exact test, p=0.099)). **B)** Pairwise tests for differentially methylated loci (DML) between *O. melanoleuca* (mel), *O. pleschanka* (ple) and hybrids (hyb) show that most DMLs are found between parental species (speciesDMLs). Between parental species and hybrids, more DMLs are shared between mel vs. ple and mel vs. hyb comparisons than between mel vs. ple and mel vs. hyb indicating a greater number of conserved methylation states between *O. pleschanka* and hybrids.

Even in the absence of strong hybrid-specific methylation differences, methylation states in hybrids can provide valuable insight into the evolutionary maintenance of methylation. Because hybrids carry genomic contributions from both parental species, their methylation profiles can be compared to each parental population, analogous to inheritance-mode analyses developed for gene expression data (Wittkopp et al. 2004; McManus et al. 2010). For each CpG site, we quantified the difference in mean methylation levels between hybrids and each parental population. Joint consideration of these two hybrid-parent contrasts enabled classification of CpG sites as additive (hybrid intermediate between parents), dominant (hybrid matching one parent), transgressive (hybrid outside parental range), or conserved (difference <20% relative to both parents).

To identify sites suitable for inheritance analysis, we first screened methylomes for population differences using a generalized linear mixed model implemented in DSS, with population assignment as predictor and genome-wide mean methylation as covariate. This analysis identified 1,226 CpG sites showing significant methylation differences among populations (FDR < 0.05).

When all hybrids were analyzed as a single group, the majority of these CpG sites (n = 748; 60.9%) were classified as conserved, showing less than 20% methylation difference relative to both parental species (**fig. 5A**). Among sites exceeding this threshold, 104 (10.6%) were classified as additive, whereas 401 (32.7%) showed dominance toward one parental species. Dominance was asymmetric: 254 CpG sites were *O. pleschanka-*dominant and 147 were *O. melanoleuca-*dominant (**fig. 5E**). In this analysis, no site showed a transgressive signature.

**Figure 5.**
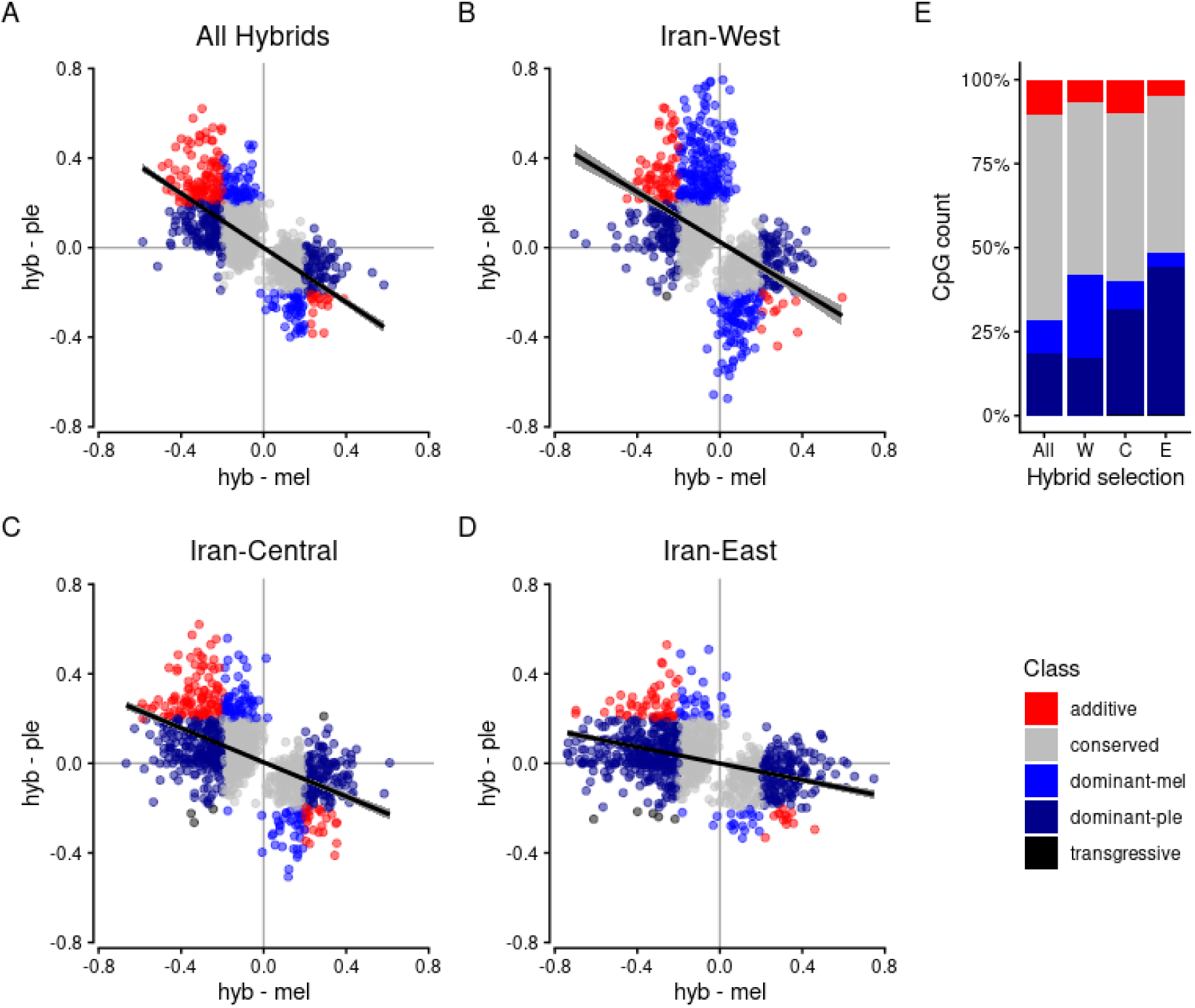
Similarity of methylation levels in hybrid subpopulations follows geographical and genetic proximity to parental populations. ‘Inheritance’-style plots for **A)** all Iranian hybrid, and hybrid subpopulations in **B)** western Iran, **C)** central Iran, and **D)** eastern Iran. Following the admixture gradient from *O. melanoleuca* (mel, west) and *O. pleschanka* (ple, east), the amount of methylation states shifts from more similar to *melanoleuca* (dominant-mel) to *pleschanka* (dominant-ple). Note that central Iran shows a higher amount of *pleschanka*-dominant CpG site, indicating asymmetry in the hybrid zone. **E)** Proportions of inheritance classes for each hybrid subpopulation

To better understand how ancestry proportions contribute to determining methylation levels across the hybrid zone, we repeated the analysis with hybrids not treated as a single population, but divided into populations from western, central, and eastern Iran. To this end we used genetically determined assignments into three hybrid subpopulations (see Materials and Methods). Unbiased hierarchical clustering of methylation levels at differentiated CpG sites recapitulated this pattern (**supplementary fig. S9**, Lutgen et al. 2025), thus reaffirming using these assignments for analyses of methylation variation and providing another indication for strong association between genetic and methylation variation.

Based on this framework, we quantified methylation patterns in hybrids across the Iranian hybrid zone as well as within hybrid populations from western, central, and eastern Iran, which exhibit different proportions of genetic ancestry from *O. melanoleuca* and *O. pleschanka*. Western and eastern hybrids show higher ancestry proportions of the neighboring parental species than central hybrids that have more equal ancestry proportions of *O. melanoleuca* and *O. pleschanka* (**supplementary fig. S10**) Unbiased hierarchical clustering of methylation levels at differentiated CpG sites recapitulated this pattern (**supplementary fig. S9**, Lutgen et al. 2025) and thus reaffirmed using these assignments for our analyses.

Consistent with these ancestry gradients, the proportion of dominance signatures shifted longitudinally across the hybrid zone. In eastern Iran, 44.0% of classified sites were *O. pleschanka*–dominant compared to 3.8% *O. melanoleuca-*dominant. In western Iran, this contrast was reduced (24.6% vs. 17.2%). Central hybrids showed 31.2% *O. pleschanka-*dominant and 8.5% *O. melanoleuca-*dominant sites. Thus, dominance frequencies track changes in genetic ancestry proportions across the hybrid zone.

Across all hybrid subpopulations, nine CpG sites were classified as transgressive. Five occurred in eastern hybrids and four in central hybrids, with one site shared between central and western populations. Eight of these sites were located in intergenic regions or annotated repeats or repeat flanks. One transgressive CpG site occurred within the coding sequence of Znf710, a zinc finger protein predicted to function as a transcription factor.

Finally, we examined whether chromosomes exhibiting elevated parental differentiation differed in inheritance patterns. As shown previously (**fig. 3B**), chromosomes 4A and Z harbored higher numbers of loci differentially methylated between parental species. However, neither chromosome contained transgressive sites. Instead, they differed in the relative frequency of dominance signatures. On chromosome Z, *O. pleschanka*-dominant sites were more frequent than *O. melanoleuca-*dominant sites (44 vs 3), whereas this pattern was reversed on chromosome 4A. Additive sites were relatively more abundant on chromosome 4A than on chromosome Z (**supplementary fig. S11**).

In summary, the abundance of parental-dominant methylation signatures shifts longitudinally across the hybrid zone, paralleling changes in genetic ancestry from more *melanoleuca*-like in the west to more *pleschanka*-like in the east. With only nine of 1,226 CpG sites showing methylation levels outside the parental range, evidence for transgressive methylation is rare. Although chromosomes 4A and Z exhibit elevated parental differentiation, neither harbors transgressive sites. Instead, they differ in the relative frequency of *O. pleschanka*- and *O. melanoleuca*-dominant CpG sites, indicating chromosome-specific differences in methylation inheritance patterns.

## Discussion

By disrupting methylation, hybridization can induce a molecular cascade that may reduce the fitness of hybrids (Brown et al. 2012; Lauss et al. 2018; Kakoulidou et al. 2024), and, ultimately, contribute to reproductive barriers (Smith et al. 2016; Laporte et al. 2019; Lammers et al. 2026). For such a scenario to unfold, we expect methylation patterns in hybrids to differ from the ones observed in parental species, because co-evolved regulators of methylation become uncoupled from their target location in admixed genomes. To understand how methylation patterns can change under natural hybridization, we here investigated the variation of methylation in a natural avian hybrid zone. Based on 100 methylomes from two closely related species, we find that across more than 436,000 CpG sites methylation variation closely mirrors genetic variation. Notably, population structure is highly similar between methylation and genetic variation, and, within hybrids, genome-wide genetic ancestry predicts a substantial proportion of methylation variation. Methylation patterns of parental species differ at a rather limited number of CpG sites, and these differences largely co-localize with elevated genetic differentiation at the level of whole chromosomes and to a lesser degree at the level of location along chromosomes. Finally, we find no evidence for substantial hybrid-specific, respectively, transgressive methylation patterns that would be indicative of widespread disrupted methylation regulation in hybrids. Together, these results indicate that methylation in this system is primarily shaped by underlying genetic variation, even in the context of extensive hybridization.

Our results contribute to a growing body of work suggesting that genetic control of methylation variation is widespread in the animal kingdom. This is reflected by the correlation of both the geographic structure of methylation and genetic variation, and of methylation divergence with genetic differentiation between species. First, across the *hispanica* complex, genome-wide genetic ancestry emerges as a robust predictor of methylation variation (**fig. 2**) Specifically, the population structure of methylation confidently recapitulates genetic population structure, with the similarity of methylation among individuals as inferred by PCA closely matching individuals’ genetic similarity inferred from genome-wide SNP data (**fig. 2**). This correspondence between methylation and genetic variation is particularly striking in hybrids, where a substantial fraction of methylation variation is almost perfectly predicted by genetic variation (**fig. 2C**). Similar correlations between genetic and methylation variation are documented from various animals, including humans (Carja et al. 2017), great apes (Hernando-Herraez et al. 2015), and stick insects (de Carvalho et al. 2023). Within birds, our results are consistent with studies in crows (Merondun and Wolf 2025), great tits (Sepers, Chen, et al. 2023), and flycatchers (Boman et al. 2024) that also report a pronounced genetic determination of methylation. Our work extends these findings from species with hybridization limited to a single generation (flycatchers; Kawakami et al. 2014) or near to absent genome-wide differentiation of hybridizing population (crows; Poelstra et al. 2014) with population-scale insights from a genetically well-differentiated and at the same time pervasively admixed system (Schweizer et al. 2019; Alaei Kakhki et al. 2023; Lutgen et al. 2025).

A second line of evidence supporting an important role of genetics in the regulation of methylation, which, moreover, may provide insights into the evolutionary and molecular processes underlying this genetic regulation stems from the correlation of methylation divergence and genetic differentiation between *O. melanoleuca* and *O. pleschanka*. The methylation divergence between species that is observed for each chromosome strongly correlates with chromosomes’ genetic differentiation (**fig. 3C**). Within this correlation, the two chromosomes with significant enrichment of DMLs, chromosomes Z and 4A, are also genetically most differentiated (**fig. 3C**). For chromosome Z, this coincidence of methylation divergence and genetic differentiation may be explained by its function as a sex chromosome. The high genetic differentiation is expected, owing to the smaller effective population size relative to autosomes, as has been widely documented (e.g. Ellegren et al. 2012; Wang et al. 2014; Irwin 2018; Hayes et al. 2020). For methylation divergence, we can only speculate that the high methylation divergence we observe on chromosome Z may be linked to fast divergence of male-hypermethylation in restricted regions of chromosome Z (Höglund et al. 2024). In contrast, chromosome 4A may reflect correlations between methylation and genetics documented for autosomes of other species (Hernando-Herraez et al. 2015; de Carvalho et al. 2023; Ord et al. 2023; de Carvalho et al. 2025), where the correlation often is not limited to chromosome-wide averages, but methylation divergence and genetic differentiation correlated along chromosomes. These correlations have alternatively been explained by the hypermutability of methylated CpGs (Holliday and Grigg 1993) or modified regulation of methylation by sequence variation in genes encoding for methylation-regulating proteins (see also next paragraph). In the former scenario, methylation precedes genetic differentiation because methylation leads to a deamination of methylated cytosines and consequently elevated mutation rates (Smith et al. 2016; Ord et al. 2023). Alternatively, sequence divergence may directly cause methylation differences by regulating methylation also in trans, for instance via altered transcription factor binding or chromatin organization (Hawe et al. 2022; Paul 2025). Under this model sequence evolution controls methylation divergence also at distant loci. Because our results show only partial co-localization of DMLs with highly differentiated regions on chromosomes (**fig. 3D, E**; **supplementary fig. S6**), we interpret our results to be mostly consistent with a scenario in which methylation follows genetic differentiation rather than preceding it.

Our evidence for a genetic component do not rule out environmental contributions to methylation variation, both of which are well documented (Anastasiadi et al. 2017; Sheldon et al. 2020; Sepers et al. 2021). Still, it would seem unlikely that environmental variation would be driving the correlation of geographic structures of methylation and genetic variation observed here and elsewhere (Hernando-Herraez et al. 2015; Carja et al. 2017; de Carvalho et al. 2025). In such a case, we would not expect the nearly perfect correlation of population structures (**fig. 2C**), especially since in the centre of the hybrid zone individuals from a broad admixture spectrum are exposed to highly similar environments (Lutgen et al. 2025). Therefore, irrespective of potential contributions of the environment or life history challenges to methylation regulation (Sheldon et al. 2020; Lauer et al. 2023), our results add to a growing body of evidence that methylation is strongly constrained, if not largely determined, by the genetic background. Next, the correlation of methylation divergence with genetic differentiation along chromosomes, such as for instance observed for chromosome 4A, may provide a more nuanced perspective on the trans versus cis regulation of methylation that suggests how different regulatory architectures of methylation may participate in methylation regulation across the genome. Previous studies reported considerable variation in the abundance of cis- and trans-associated meQTLs. In hybridizing carrion and hooded crows, CpG methylation appears to be mostly cis-regulated (Merondun and Wolf 2025). Meanwhile, in pied and collared flycatcher proportions of cis versus trans regulated methylation were more balanced, depending on tissue (Boman et al. 2024). In contrast to this, but similar to an intraspecific study in great tits (Sepers, Chen, et al. 2023), out of the 150,000 associations between genetic variants and methylation levels identified here, 88.3% are trans-associated. Although meQTL analyses may be have limited power due to rather low sample sizes or model fit (Mueller et al. 2025), the predominance of trans-associated meQTLs reported here is within the range of other avian studies and consistent with a regulatory architecture in which distal genetic factors play a major role in determining methylation states. Notably, the distinct co-localization of (i) DMLs with high *F*_ST_ regions and (ii) a more balanced distribution of DMLs along chromosomes show that cis and trans regulation of methylation may vary across the genome. Together with the abovementioned contrasting results between systems, this indicates that the regulatory architecture of methylation may differ considerably within and across species. Controlled hybridization experiments incorporating broader taxonomic sampling across multiple tissues, together with analyses of allele-specific methylation, will enable a more detailed disentanglement of the regulatory architecture underlying methylation.

Finally, we find little evidence for transgressive methylation in wheatear hybrids, indicating that hybridization in this system does not broadly disrupt methylation regulation. Despite an important genetic component in the regulation of methylation, methylation levels may become transgressive in hybrids (Berbel-Filho et al. 2022), that is be outside methylation levels observed in parental species. In contrast to hybridization-induced transgressive methylation patterns, such as observed in *Larix* (Li et al. 2013), *Arabidopsis* (Lauss et al. 2018) or *Coregonus* (Laporte et al. 2019), methylation levels in hybrid wheatears show a stable transmission of methylation, that is, hybrids’ methylation levels come to lie between parental methylation levels (**fig. 4B**) and are well predicted by hybrids’ ancestry composition (**fig. 5**). Only eight CpG sites (out of 1,226) exhibited transgressive methylation (**fig. 5**; **supplementary table S5**). This indicates that hybridization in the studied wheatears does not lead to a systemic breakdown of methylation regulation. Instead, hybrid methylation levels largely fall within the parental range of variation and are characterized predominantly by additive or dominant transmission patterns. Notably, dominance patterns shift toward the genetically closer parental species at the western and eastern edges of the hybrid zone (**fig. 5C–E**). This reinforces the conclusion that methylation variation in hybrids closely tracks genome-wide ancestry. Together, these results provide little support for hybridization-induced instability of methylation.

This result does not fit predictions made by the concept of hybridization-induced “genomic shock,” originally proposed in plants (McClintock 1984) and later supported by empirical evidence in vertebrate systems (Dion-Côté et al. 2014; Laporte et al. 2019). The “genomic shock” hypothesis postulates that the repression of transposable element (TE) activity is disrupted upon hybridization when the co-inheritance of TEs and their repressors is broken up. Because methylation is a primary repressor of TE activity in animals and plants, transgressive methylation is expected to be a key signature of this disruption. Different from previous studies that found broad TE demethylation in hybrids (Brown et al. 2012; Laporte et al. 2019), we detected only two transgressively methylated sites within TEs, of which neither is associated with a TE retaining clear functional potential. Several explanations may account for the stability of hybrid methylation in wheatears. First, avian methylomes may be comparatively robust to genetic and environmental perturbation, as suggested by population-level studies in birds (Sepers, Chen, et al. 2023; Sepers, Mateman, et al. 2023; Merondun and Wolf 2025), but see Sheldon et al. 2020) for possible environmental influence. Second, a disruption of methylation regulation may happen predominantly during earlier generations of hybridization, and, notably, backcrossing but be resolved in old hybrid zones. The hybrid zone in wheatears studied here may represent such an older system that moved already past a broad destabilization of methylation regulation, where variation underpinning disrupted methylation was already purged or stabilized over time. From this perspective, our results would, if methylation was ever disrupted, support a scenario in which hybrids recovered from initial hybrid-induced breakdown by evolving compensatory regulatory architectures that stabilized methylation in later generation hybrids, as was shown for methylation in hybrid flycatchers (Boman et al. 2024) or gene misexpression in hybrids of *Drosophila* (Wittkopp et al. 2004). Because the Iranian hybrid zone between *O. melanoleuca* and *O. pleschanka* studied here consists merely of late-generation and no early-generation hybrids, this hypothesis cannot be directly tested. Third, it is also possible that the low genetic divergence between *O. melanoleuca* and *O. pleschanka* is insufficient to trigger substantial disruption of methylation maintenance. Additional studies in systems with greater genetic differentiation or younger hybrid zones, and, ideally, controlled crosses under experimental conditions, are called for to answer these questions.

## Conclusion

We conclude that methylation variation in the *O. hispanica* complex is tightly associated with genetic ancestry, exhibits limited divergence between species, and remains largely stable in hybrids. Rather than acting as an independent or rapidly evolving regulatory layer, methylation in this system primarily reflects underlying genomic differentiation and is robust to genome reshuffling through hybridization. Together, these findings suggest that methylation is unlikely to constitute a major driver of reproductive isolation or hybrid dysfunction in this system but instead evolves largely as a downstream consequence of genetic divergence.

## Materials and Methods

### CpG annotation of the reference genome

Our analysis used the chromosome-level reference genome of *O. melanoleuca* (OenMel1.1) and its annotation of genes and repetitive elements (Peona et al. 2023, Lutgen et al. 2025). We extended this annotation by identifying CpG islands (CGIs) using makeCGI v.1.34 (Wu et al. 2010). We filtered predicted CGIs to retain regions at least 200 bp in length, with a GC content of at least 50% and an observed-to-expected CpG ratio of at least 0.6; the resulting set constituted the final CGI annotation.

To assign each CpG dinucleotide a unique genomic context, we extracted CpG coordinates from the reference genome and applied a hierarchical annotation strategy (Merondun and Wolf 2025). We classified CpG sites, in decreasing order of priority, as belonging to promoter-associated CGIs (CGIs 2000 bp upstream of genes), non-promoter CGIs, repetitive elements, repeat flanking regions, coding sequences, introns, or intergenic regions. This prioritization ensured unambiguous assignments while favoring CpG sites with higher presumed functional relevance. We stored all CpG annotations in a custom BED-like format compatible with downstream processing using the bedtools suite (Quinlan and Hall 2010).

### Sampling and Sequencing

To generate whole-genome methylation data for the *hispanica* complex, we sequenced 101 individuals representing four species (Schweizer et al., 2019a; **supplementary table 1**). We extracted genomic DNA from ethanol-preserved blood samples of western black-eared wheatears (*O. hispanica*), pied wheatears (*O. pleschanka*), eastern black-eared wheatears (*O. melanoleuca*), and Cyprus wheatears (*O. cypriaca*) using the MagAttract HMW DNA Kit (Qiagen) following the manufacturer’s protocol.

To prepare sequencing libraries, we performed enzymatic conversion-based library preparation using the NEBNext® Enzymatic Methyl-seq Kit (New England Biolabs), following the manufacturer’s protocol, including a spike-in of phage lambda to estimate conversion rate of unmethylated cytosines (see below). All libraries were sequenced on an Illumina NovaSeq 6000 instrument with target coverage 15x.

### Individualized, SNP-corrected reference genomes

To account for C>T (and G>A) substitutions that are indistinguishable from cytosine conversion artifacts introduced during enzymatic methyl-seq (EM-seq), and to improve alignment efficiency, we generated individualized, SNP-corrected reference genomes for each sample using published whole-genome sequencing (WGS) data from the same individuals (Lutgen et al. 2025). We extracted high-quality single-nucleotide polymorphisms (SNPs) from WGS reads mapped to the OenMel1.1 reference genome (Peona et al. 2023) and incorporated these variants into consensus sequences using bcftools consensus (Danecek and McCarthy 2017). SNPs were filtered by excluding variants with evidence of strand bias (FS > 60.0 or SOR > 3.0), low variant confidence relative to depth (QD < 2.0), poor mapping quality (MQ < 40.0), or significant mapping- or position-based biases (MQRankSum < −12.5 or ReadPosRankSum < −8.0). To preserve coordinate consistency across all reference genomes, we excluded insertions and deletions from the consensus generation and transferred SNPs only. For one sample (GR-LES-2017-001) lacking a corresponding WGS dataset, we used a genetically close individual (GR-LES-2017-002) as a surrogate for SNP correction. We validated all SNP-corrected genomes by checking sequence length and base composition for consistency with the original reference. To enable downstream methylation-aware alignment, we generated reference genome indices using the Bismark genome preparation tool (Krueger and Andrews 2011).

### Read mapping and processing of EMSeq data

To prepare EM-seq reads for alignment, we trimmed sequencing adapters and low-quality bases using trim_galore 0.6.10 (Krueger et al. 2023) with settings optimized for enzymatic methylation sequencing (--clip_r1 8, --clip_r2 8, --three_prime_clip_r1 8, --three_prime_clip_r2 8). We mapped trimmed reads to the sample-specific SNP-corrected reference genomes using Bismark v0.24.1 in combination with Bowtie2 v2.5.1 (Krueger and Andrews 2011; Langmead and Salzberg 2012) applying the parameters --score_min L,0,-0.2, --dovetail, and --maxins 1000. To remove PCR duplicates, we used the Bismark deduplication module. We then extracted methylation information using Bismark’s methylation extractor, which produced strand-specific counts of cytosine and thymine reads at each CpG site. To generate a single methylation call per CpG dinucleotide, we combined strand-specific counts using custom scripts (Lammers 2026), thereby producing destranded per-CpG methylation readouts for downstream analyses. Cytosine conversion rates were estimated by mapping the libraries to phage lambda using the same parameters and were between 96.6% and 100.0% (mean 99.5±0.5%).

### Processing of methylation calls

Destranded methylation calls obtained from Bismark were processed with the R-package methylKit v0.99.2 (Akalin et al. 2012). We filtered CpG sites to remove extreme coverage outliers by excluding sites above the 99.5^th^ percentile of coverage. For all analyses, except those performed with DSS that include coverage estimates in modeling methylation differences (see below), we required a minimum coverage of five reads per CpG site. To account for differences in sequencing depth across samples, we normalized coverage using scaling factors calculated from between-sample differences in median coverage, as implemented in methylKit. After visually inspecting coverage distributions and methylation profiles, we excluded three samples due to low mapping rate. We then merged (methyKit unite) methylation data across the remaining 98 samples, retaining 436,762 CpG sites with complete methylation information for downstream analyses.

### Principal component analysis (PCA) of methylation

To assess the population structure of methylation variation, we performed PCA on CpG methylation profiles derived from the methylKit “unite” tables. For each CpG site in each sample, methylation proportion was calculated as

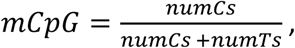

where numCs is the number of methylated Cs and numTs is the number of unmethylated Cs (sequenced as T after enzymatic conversion). Sites were identified by genomic coordinate (chromosome:position), and data were reshaped to a matrix with samples as rows and CpG sites as columns. PCA was computed with the R-package pcaMethods 2.2.0 (Stacklies et al. 2007), retaining the first 20 principal components (PCs). To visualize the major population splits between the focal populations *O. melanoleuca, O. pleschanka*, their hybrid zone, as well as the Pontic lineage and *O. hispanca* in a single PCA plot we combined PC3 and PC4, which separated these lineages best, by summing over the two PC axes.

To investigate the geographic structure of methylation variation, we modeled the relationship between longitude and PC2. We chose PC2 because it captures most variation that separates the focal lineages, *O. melanoleuca* and *O. pleschanka*, and thus was most likely to reflect the geographic structure. To this end, we modeled spatial variation in PC2 using a four-parameter logistic cline as follows:

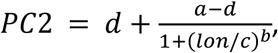

where a and d represent the maximum and minimum of the PC2 range (extremities of the hybrid zone), c is the mean of PC2 (inflection point), b controls the slope (b=1), and lon is longitude. This model equivalent to the classical hybrid-zone models described by Barton and Gale (1993) and used to describe geographic transitions in allele frequencies and quantitative traits (Stankowski 2013).

The model was fitted using nonlinear least squares with the Levenberg-Marquardt algorithm (nlsLM() from the minpack.lm R package). Starting values were derived from summary statistics of the data. Confidence intervals for model predictions were obtained via error propagation using predictNLS() from the propagate package. Model fit was evaluated using a pseudo-R^2^ (calculated as *R*^2^ = 1 − *RSS/TSS*, with RSS=residual sum of squares, and TSS=total sum of squares).

PCA results for genetic data were obtained from published analyses (Lutgen et al. 2025). In brief, genotype frequencies were calculated from strictly filtered SNPs, and PCA was performed using smartpca (Patterson et al. 2006) with a stepwise approach to ensure balanced sample composition between lineages (Lutgen et al. 2025).

### Calling of differentially methylated sites and regions

To identify CpG sites exhibiting significant methylation differences among populations, we used the statistical framework implemented in DSS 2.49.0 (Feng and Wu 2019), which models counts of methylated and unmethylated cytosines as beta-binomial distribution and tests for differentially methylated sites (DMLs) and differentially methylated regions (DMRs) using a dispersion shrinkage method and Wald test. We applied a two-group pairwise comparison for methylation differences between *O. melanoleuca* and *O. pleschanka.* DMLs and DMRs were called using an initial p-value threshold of 0.001 and after Benjamin-Hochberg correction for multiple testing, filtered for FDR < 0.05 and an absolute methylation difference ≥0.2. To generate methylation differences at these inter-species DMLs, we plotted the mean methylation level per site and species as returned by DSS, using ComplexHeatmap (Gu et al. 2016). To generate a clustering dendrogram of similar methylation differences between both populations, we calculated euclidean distances as implemented in R’s dist() function that were then clustered using the ward.D2 algorithm as implemented (’hclust()’ function).

To generate the Venn diagram of DMLs shared between parental species and hybrids we aimed at identifying overlapping DMLs between all three populations (fig. 4B). First, to call DMLs in pairwise comparisons across these three populations, we included only samples that were unambiguously assigned to *O. melanoleuca*, *O. pleschanka*, or the hybrid population using genetically determined population assignments from a published FineStructure matrix (Lutgen et al. 2025). DMLs were called using DSS as described above and filtered using the parameters as before. Then, to obtain a unified list of differentially methylated CpG sites across all three populations, we combined the DMLs called in the three pairwise comparisons. From this, we counted overlapping DML coordinates between the three pairwise comparisons to build the Venn diagram of with the R package ggVennDiagram version 1.5.7 (Gao C, Dusa A (2026)).

### Test for chromosomal enrichment of DMLs

To test whether DMLs were non-randomly distributed across chromosomes, we compared the observed number of DMLs per chromosome to expectations based on the number of CpG sites tested per chromosome. Expected counts were calculated by scaling chromosome-specific CpG counts to the total number of detected DMLs. A χ^2^ goodness-of-fit test was used to assess the overall deviation from the expected distribution. In addition, chromosome-specific enrichment or depletion of DMLs was evaluated using binomial tests, with p-values adjusted for multiple testing using the Benjamini–Hochberg procedure. Enrichment was quantified as the log_2_-ratio of observed to expected counts.

### Methylation QTL (meQTL) analysis

To identify genetic variants associated with methylation levels (meQTLs), we used the MatrixEQTL R package (Shabalin 2012), adapted for meQTL mapping. Genotype data for matching individuals were extracted from a multi-sample VCF file available from Lutgen et al. (2025) and converted to dosage format per SNP and sample, with dosage being the number of alternative alleles (0, 1, 2). Methylation data were provided as a matrix of methylation levels per CpG.

To include only high-quality SNPs in the analysis, we filtered for biallelic SNPs with a minor allele frequency (MAF) ≥ 0.05 and ≤ 0.95 and missingness of ≤ 0.05. CpG sites with missing values or low variability across samples were excluded. We defined cis-meQTL pairs as SNP-CpG pairs located within 1 Mb of each other (Shabalin 2012. Sepers et al, 2023).

meQTL analysis was performed using linear models of genotypes as predicting variable and the first four genetic principal components and mean sequencing depth as covariates to account for population structure and technical variability, respectively. Significance was assessed using the built-in t-statistic, and multiple testing correction was applied using the Benjamini–Hochberg false discovery rate (FDR) method. Only associations with FDR-adjusted p-values below 0.05 were considered significant.

### Identification of methylation transmission modes

To visualize how methylation level in hybrids compare to those in the parental populations, we adopted inheritance-mode analyses developed for gene expression data (Wittkopp et al. 2004; McManus et al. 2010) that are based on plotting differences of methylation levels between hybrids to both parental populations. To this end, we first screened for differentially methylated sites across all three populations using a generalized linear model implemented in DSS (DMLfit.multiFactor()) (Park and Wu 2016). The model uses an arcsine link function to accommodate methylation proportions that approach the boundaries of 0 and 1. Linear model fitting is then performed using ordinary least squares on the transformed methylation levels. In this model, population assignment was included as a predictor variable, enabling us to test for differences among groups. We also included the first principal component from the PCA (see above) as covariate to account for additional sources of variance (mostly interindividual differences in genome-wide methylation levels) captured by this axis. From the fitted model, we extracted the test statistics for the coefficients given by the three population-contrasts (i.e. *melanoleuca* vs. *pleschanka*, *melanoleuca* vs. hybrids, *pleschanka* vs. hybrids) using DMLtest.multiFactor(). To get a unified list of differentially methylated sites, we combined the test statistics, recalculated p-values using Bonferroni correction if the same CpG site was returned for multiple contrasts. Finally, false discovery rates were calculated using the Benjamini–Hochberg procedure, and CpG sites were filtered using an FDR threshold of 0.05.

For each CpG site, we quantified the difference in mean methylation levels between hybrids and each parental population. Joint consideration of these two hybrid–parent contrasts enabled classification of CpG sites as additive (hybrid intermediate between parents), dominant (hybrid matching one parent), transgressive (hybrid outside parental range), or conserved (difference <20% relative to both parents).

These contrasts enable placing hybrid methylation states within a two-dimensional space, whose axes describe their (dis-)similarity to either parental population and make it possible to differentiate between modes of transmission that can be ‘additive’ (i.e. intermediate state between parental species), ‘dominant’ (i.e. biased towards one parental state) and ‘transgressive’ (i.e. exceeding both parental states). To describe these different signatures of methylation transmission in hybrids, we adopt terminology from gene expression studies that infer inheritance modes by contrasting expression levels between parental and hybrid transcriptomes. However, here we use these different inheritance modes to describe patterns of methylation transmission established over evolutionary timescales, not the immediate outcome of transgenerational inheritance in parent –offspring comparison that is used in experimental studies of early-generation hybrids.

### Clustering of pairwise similarities of methylation variation

To visualize pairwise similarities between genome-wide methylation levels across samples, we generated a heatmap of pairwise distances between samples generated from correlation matrices. Correlation matrices of methylation levels were calculated with R’s cor() function using Spearman’s ρ. Pairwise distances were then plotted as heatmaps produced with ComplexHeatmap (Gu et al. 2016) with the ward.D2 clustering method and transforming correlations to distances (1-x). All computations were performed in R 4.4.3.

## Data availability

EMSeq data generated for this study has been deposited to the SRA under BioProject PRJNA1134926. All whole-genome sequencing data was published earlier under BioProject PRJNA1128009.

## Acknowledgements

We acknowledge funding from the Swiss National Science Foundation (SNSF) through grant 310030_207417 to R.B. and a Swedish Research Council (Vetenskapsrådet) grant 2022-06195 to V.P. We thank Pius Korner for statistical support, Sarah Mueller and the Wolf group, and Manuel Schweizer for discussion. We thank the Next Generation Sequencing Platform of the University of Bern for performing the high-throughput sequencing experiments. Computations were performed at the Genetic Diversity Centre (GDC), ETH Zurich.

## References

Abbott R, Albach D, Ansell S, Arntzen JW, Baird SJE, Bierne N, Boughman J, Brelsford A, Buerkle C a., Buggs R, et al. 2013. Hybridization and speciation. J. Evol. Biol. 26:229–246.

Akalin A, Kormaksson M, Li S, Garrett-Bakelman FE, Figueroa ME, Melnick A, Mason CE. 2012. methylKit: a comprehensive R package for the analysis of genome-wide DNA methylation profiles. Genome Biol. 13:R87.

Al Adhami H, Bardet AF, Dumas M, Cleroux E, Guibert S, Fauque P, Acloque H, Weber M. 2022. A comparative methylome analysis reveals conservation and divergence of DNA methylation patterns and functions in vertebrates. BMC Biol. 20:70.

Anastasiadi D, Díaz N, Piferrer F. 2017. Small ocean temperature increases elicit stage-dependent changes in DNA methylation and gene expression in a fish, the European sea bass. Sci. Rep. 7:12401.

Berbel-Filho WM, Pacheco G, Lira MG, Garcia de Leaniz C, Lima SMQ, Rodríguez-López CM, Zhou J, Consuegra S. 2022. Additive and non-additive epigenetic signatures of natural hybridization between fish species with different mating systems: Running head: Epigenetics of fish hybrids. Epigenetics 17:2356–2365.

Bird A. 2002. DNA methylation patterns and epigenetic memory. Genes Dev. 16:6–21.

Boman J, Qvarnström A, Mugal CF. 2024. Regulatory and evolutionary impact of DNA methylation in two songbird species and their naturally occurring F1 hybrids. BMC Biol. 22:124.

Brown JD, Piccuillo V, O’Neill RJ. 2012. Retroelement demethylation associated with abnormal placentation in Mus musculus x Mus caroli hybrids. Biol. Reprod. 86:88.

Carja O, MacIsaac JL, Mah SM, Henn BM, Kobor MS, Feldman MW, Fraser HB. 2017. Worldwide patterns of human epigenetic variation. Nat. Ecol. Evol. 1:1577–1583.

de Carvalho CF, Planidin NP, Villoutreix R, Soria-Carrasco V, Riesch R, Feder JL, Slate J, Nosil P, Gompert Z. 2025. Linking DNA methylation to localised genetic differentiation in Timema cristinae stick insects. Mol. Ecol. 34:e70049.

de Carvalho CF, Slate J, Villoutreix R, Soria-Carrasco V, Riesch R, Feder JL, Gompert Z, Nosil P. 2023. DNA methylation differences between stick insect ecotypes. Mol. Ecol. 32:6809–6823.

Chapelle V, Silvestre F. 2022. Population epigenetics: The extent of DNA methylation variation in wild animal populations. Epigenomes 6:31.

Danecek P, McCarthy SA. 2017. BCFtools/csq: haplotype-aware variant consequences. Bioinformatics 33:2037–2039.

Dion-Côté A-M, Renaut S, Normandeau E, Bernatchez L. 2014. RNA-seq Reveals Transcriptomic Shock Involving Transposable Elements Reactivation in Hybrids of Young Lake Whitefish Species. Mol. Biol. Evol. 31:1188–1199.

Feng H, Wu H. 2019. Differential methylation analysis for bisulfite sequencing using DSS. Quant. Biol. 7:327–334.

Goncalves A, Leigh-Brown S, Thybert D, Stefflova K, Turro E, Flicek P, Brazma A, Odom DT, Marioni JC. 2012. Extensive compensatory cis-trans regulation in the evolution of mouse gene expression. Genome Res. 22:2376–2384.

Gu Z, Eils R, Schlesner M. 2016. Complex heatmaps reveal patterns and correlations in multidimensional genomic data. Bioinformatics 32:2847–2849.

Hawe JS, Wilson R, Schmid KT, Zhou L, Lakshmanan LN, Lehne BC, Kühnel B, Scott WR, Wielscher M, Yew YW, et al. 2022. Genetic variation influencing DNA methylation provides insights into molecular mechanisms regulating genomic function. Nat. Genet. 54:18–29.

Hernando-Herraez I, Heyn H, Fernandez-Callejo M, Vidal E, Fernandez-Bellon H, Prado-Martinez J, Sharp AJ, Esteller M, Marques-Bonet T. 2015. The interplay between DNA methylation and sequence divergence in recent human evolution. Nucleic Acids Res. 43:8204–8214.

Holliday R, Grigg GW. 1993. DNA methylation and mutation. Mutat. Res. 285:61–67.

Kakoulidou I, Johannes F. 2024. DNA methylation remodeling in F1 hybrids. Plant J. 118:671–681.

Kakoulidou I, Piecyk RS, Meyer RC, Kuhlmann M, Gutjahr C, Altmann T, Johannes F. 2024. Mapping parental DMRs predictive of local and distal methylome remodeling in epigenetic F1 hybrids. Life Sci. Alliance [Internet] 7. Available from: 10.26508/lsa.202402599

Krueger F, Andrews SR. 2011. Bismark: a flexible aligner and methylation caller for Bisulfite-Seq applications. Bioinformatics 27:1571–1572.

Krueger F, James F, Ewels P, Afyounian E, Weinstein M, Schuster-Boeckler B, Hulselmans G, sclamons. 2023. FelixKrueger/TrimGalore: v0.6.10. Zenodo Available from: 10.5281/ZENODO.7598955

Lammers F. 2026. fritjoflammers/bioscripts: v2025.1. Zenodo Available from: 10.5281/zenodo.18495433

Lammers F, Peona V, Burri R. 2026. Transposable elements as drivers of reproductive isolation: A framework for testing hybridization-induced escalation of genetic conflicts. EcoEvoRxiv.

Langmead B, Salzberg SL. 2012. Fast gapped-read alignment with Bowtie 2. Nat. Methods 9:357–359.

Laporte M, Le Luyer J, Rougeux C, Dion-Côté A-M, Krick M, Bernatchez L. 2019. DNA methylation reprogramming, TE derepression, and postzygotic isolation of nascent animal species. Sci. Adv. 5:eaaw1644.

Lauer ME, Kodak H, Albayrak T, Lima MR, Ray D, Simpson-Wade E, Tevs DR, Sheldon EL, Martin LB, Schrey AW. 2023. Introduced house sparrows (*Passer domesticus*) have greater variation in DNA methylation than native house sparrows. J. Hered.:esad067.

Lauss K, Wardenaar R, Oka R, van Hulten MHA, Guryev V, Keurentjes JJB, Stam M, Johannes F. 2018. Parental DNA methylation states are associated with heterosis in epigenetic hybrids. Plant Physiol. 176:1627–1645.

Li A, Song W-Q, Chen C-B, Zhou Y-N, Qi L-W, Wang C-G. 2013. DNA methylation status is associated with the formation of heterosis in Larix kaempferi intraspecific hybrids. Mol. Breed. 31:463–475.

Lutgen D, Peona V, Chase MA, Kakhki NA, Lammers F, de Souza SG, Ducrest A-L, Burri M, Andriopoulos P, Lukhele SM, et al. 2025. A mosaic of modular variation at a single gene underpins convergent plumage coloration. Science 390:eado8005.

Mack KL, Nachman MW. 2017. Gene regulation and speciation. Trends Genet. 33:68–80.

McClintock B. 1984. The significance of responses of the genome to challenge. Science 226:792–801.

McManus CJ, Coolon JD, Duff MO, Eipper-Mains J, Graveley BR, Wittkopp PJ. 2010. Regulatory divergence in Drosophila revealed by mRNA-seq. Genome Res. 20:816–825.

McRae AF, Powell JE, Henders AK, Bowdler L, Hemani G, Shah S, Painter JN, Martin NG, Visscher PM, Montgomery GW. 2014. Contribution of genetic variation to transgenerational inheritance of DNA methylation. Genome Biol. 15:R73.

Merondun J, Wolf JBW. 2025. DNA methylation reflects Cis-genetic differentiation across the European crow hybrid zone. Mol. Ecol.:e70026.

Michalak P. 2009. Epigenetic, transposon and small RNA determinants of hybrid dysfunctions. Heredity (Edinb.) 102:45–50.

Moran BM, Payne C, Langdon Q, Powell DL, Brandvain Y, Schumer M. 2021. The genomic consequences of hybridization. Elife [Internet] 10. Available from: 10.7554/eLife.69016

Moran BM, Payne CY, Powell DL, Iverson ENK, Donny AE, Banerjee SM, Langdon QK, Gunn TR, Rodriguez-Soto RA, Madero A, et al. 2024. A lethal mitonuclear incompatibility in complex I of natural hybrids. Nature 626:119–127.

Mueller SA, Merondun J, Lečić S, Wolf JBW. 2025. Epigenetic variation in light of population genetic practice. Nat. Commun. 16:1028.

O’Neill RJ, O’Neill MJ, Graves JA. 1998. Undermethylation associated with retroelement activation and chromosome remodelling in an interspecific mammalian hybrid. Nature 393:68–72.

Ord J, Gossmann TI, Adrian-Kalchhauser I. 2023. High nucleotide diversity accompanies differential DNA methylation in naturally diverging populations. Mol. Biol. Evol. 40:msad068.

Patterson N, Price AL, Reich D. 2006. Population structure and eigenanalysis. PLoS Genet. 2:e190.

Paul S. 2025. Single nucleotide polymorphisms and epigenetic crosstalk: unraveling the intricate interplay in genomic regulation. Nucleus (Calcutta) 68:505–512.

Peona V, Palacios-Gimenez OM, Lutgen D, Olsen RA, Alaei Kakhki N, Andriopoulos P, Bontzorlos V, Schweizer M, Suh A, Burri R. 2023. An annotated chromosome-scale reference genome for Eastern black-eared wheatear (Oenanthe melanoleuca). G3 (Bethesda) [Internet]. Available from: 10.1093/g3journal/jkad088

Presgraves DC. 2010. The molecular evolutionary basis of species formation. Nat. Rev. Genet. 11:175–180.

Quinlan AR, Hall IM. 2010. BEDTools: A flexible suite of utilities for comparing genomic features. Bioinformatics 26:841–842.

Runemark A, Moore EC, Larson EL. 2024. Hybridization and gene expression: Beyond differentially expressed genes. Mol. Ecol.:e17303.

Schweizer M, Warmuth V, Alaei Kakhki N, Aliabadian M, Förschler M, Shirihai H, Suh A, Burri R. 2019. Parallel plumage colour evolution and introgressive hybridization in wheatears. J. Evol. Biol. 32:100–110.

Sepers B, Chen RS, Memelink M, Verhoeven KJF, van Oers K. 2023. Variation in DNA methylation in avian nestlings is largely determined by genetic effects. Mol. Biol. Evol.:msad086.

Sepers B, Erven JAM, Gawehns F, Laine VN, van Oers K. 2021. Epigenetics and early life stress: Experimental brood size affects DNA methylation in great tits (Parus major). Front. Ecol. Evol. 9:609061.

Sepers B, Mateman AC, Gawehns F, Verhoeven KJF, van Oers K. 2023. Developmental stress does not induce genome-wide DNA methylation changes in wild great tit (Parus major) nestlings. Mol. Ecol. [Internet]. Available from: https://onlinelibrary.wiley.com/doi/abs/10.1111/mec.16973

Shabalin AA. 2012. Matrix eQTL: ultra fast eQTL analysis via large matrix operations. Bioinformatics 28:1353–1358.

Sheldon EL, Schrey AW, Hurley LL, Griffith SC. 2020. Dynamic changes in DNA methylation during postnatal development in zebra finches*Taeniopygia guttata*exposed to different temperatures. J. Avian Biol. [Internet] 51. Available from: 10.1111/jav.02294

Smith TA, Martin MD, Nguyen M, Mendelson TC. 2016. Epigenetic divergence as a potential first step in darter speciation. Mol. Ecol. 25:1883–1894.

Stacklies W, Redestig H, Scholz M, Walther D, Selbig J. 2007. pcaMethods--a bioconductor package providing PCA methods for incomplete data. Bioinformatics 23:1164–1167.

Taylor SA, Larson EL. 2019. Insights from genomes into the evolutionary importance and prevalence of hybridization in nature. *Nat*. Ecol. Evol. 3:170–177.

de Tomás C, Vicient CM. 2023. The genomic shock hypothesis: Genetic and epigenetic alterations of transposable elements after interspecific hybridization in plants. Epigenomes 8:2.

Walter M, Teissandier A, Pérez-Palacios R, Bourc’his D. 2016. An epigenetic switch ensures transposon repression upon dynamic loss of DNA methylation in embryonic stem cells. Elife [Internet] 5. Available from: 10.7554/eLife.11418

Wang X, Werren JH, Clark AG. 2016. Allele-specific transcriptome and methylome analysis reveals stable inheritance and Cis-regulation of DNA methylation in Nasonia. PLoS Biol. 14:e1002500.

Wittkopp PJ, Haerum BK, Clark AG. 2004. Evolutionary changes in cis and trans gene regulation. Nature 430:85–88.

Wolf JBW, Bayer T, Haubold B, Schilhabel M, Rosenstiel P, Tautz D. 2010. Nucleotide divergence vs. gene expression differentiation: comparative transcriptome sequencing in natural isolates from the carrion crow and its hybrid zone with the hooded crow. Mol. Ecol. 19 Suppl 1:162–175.

Wu H, Caffo B, Jaffee HA, Irizarry RA, Feinberg AP. 2010. Redefining CpG islands using hidden Markov models. Biostatistics 11:499–514.

