## Supplementary Material for "Predominantly genetic determination and stable transmission of DNA methylation in an avian hybrid zone"

to

### Supplementary Tables

**Table S1**: Sample Details (Excel, CSV)

**Table S2**: Sequencing and Methylation Calling Details (Excel, CSV)

**Table S3 | Summary statistics of meQTL analyses.** MeQTLs were modeled using a base-filtered set of SNPs (“base”) and a LD-pruned set. Multi-SNP meQTLs are defined by having associations with >1 SNPs.

| Filter | MeQTL type | N(meQTLs) | Mean SNPs/CpG | Median SNPs/CpG | Standard Deviation | Min. SNPs/CpG | Max. SNPs/CpG |
|---|---|---|---|---|---|---|---|
| base | multi-SNP | 144,717 | 20.753908 | 5 | 58.69264 | 2 | 1,191 |
| base | single-SNP | 5,966 | 1.000000 | 1 | 0.00000 | 1 | 1 |
| LD pruned | multi-SNP | 42,493 | 8.103166 | 4 | 13.21661 | 2 | 292 |
| LD pruned | single-SNP | 6,179 | 1.000000 | 1 | 0.00000 | 1 | 1 |

**Table S4 | MeQTL counts separated by association (cis, trans) for chromosomes Z and 4A.** Trans association only include intra-chromosomal meQTLs. For size-correction, trans-association with a distance larger than 19,807,785 bp (size of chromosme 4A) were excluded.

| Chromosome | uncorrected | | size-corrected | |
|---|---|---|---|---|
| | cis | trans | cis | trans |
| **chr4A** | 61 | 0 | 61 | 9 |
| **chrZ** | 216 | 95 | 216 | 32 |

**Table S5 | Transgressive CpG sites.** Table shows the coordinates of indicative of transgressive methylation levels compared to the two parental populations. Chrom, Chromosome; Pop, Population (W)est, (C)entral, (E)astern Iran); HYB-MEL, methylation differences between average methylation levels in hybrid and *O. melanoleuca;* HYB-PLE, methylation differences between average methylation levels in hybrid and *O. pleschanka;* P-Value from for the DML call; Annotation; FDR, False Discovery Rate.

| **Chrom** | **Position** | **Pop** | **HYB-MEL** | **HYB-PLE** | **P-Value** | **Genetic Annotation** | **FDR** |
|---|---|---|---|---|---|---|---|
| chr10 | 320,618 | E | -0.21738 | -0.24855 | 0.00015 | CDS; Znf710 | 0.0329 |
| chr14 | 13,259,468 | C | -0.35196 | -0.2242 | 0.00012 | Repeat flank | 0.02731 |
| chr18 | 5,507,375 | C | -0.33678 | -0.26458 | 0.00002 | Repeat flank | 0.0087 |
| chr1A | 196,51,498 | E | -0.39908 | -0.2158 | 0.00009 | Repeat; LTR | 0.02331 |
| chr2 | 24,984,805 | E | -0.61121 | -0.24926 | <0.0001 | Repeat; LINE/CR1 | 0.00002 |
| chr2 | 30,605,963 | E | -0.32347 | -0.22371 | 0.00025 | Intergenic | 0.04575 |
| chr4 | 30,196,961 | W | -0.2569 | -0.21678 | 0.00004 | Intergenic | 0.01263 |
| chr4 | 30,196,961 | C | -0.24466 | -0.20454 | 0.00004 | Intergenic | 0.01263 |
| chr4 | 63,645,882 | C | 0.29181 | 0.20979 | <0.0001 | Intergenic | 0.00208 |
| chr6 | 7,477,681 | E | -0.29735 | -0.23912 | 0.00016 | Repeat flank | 0.03358 |

### Supplementary Figures

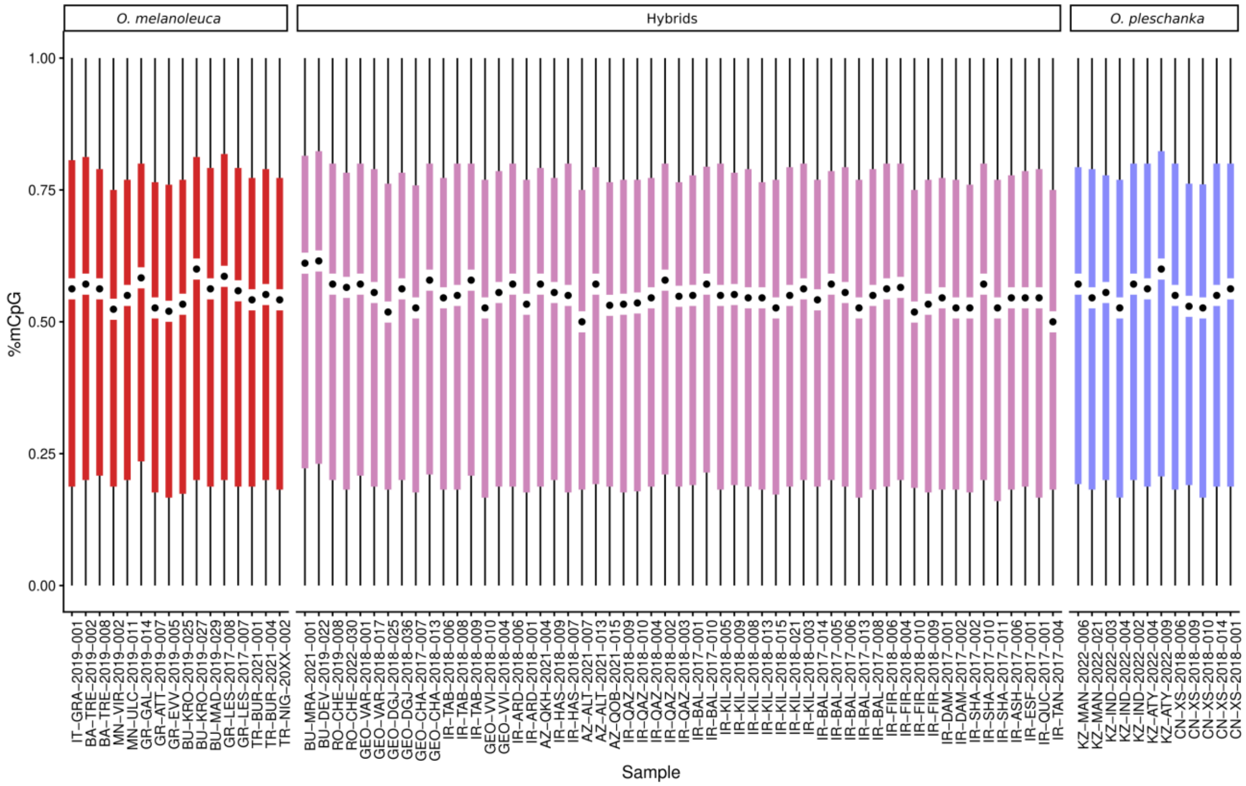


**Figure S1 | Sample-specific genome-wide methylation levels.** Per-sample boxplot of methylation levels measured as percentage of methylated Cs per CpG site.

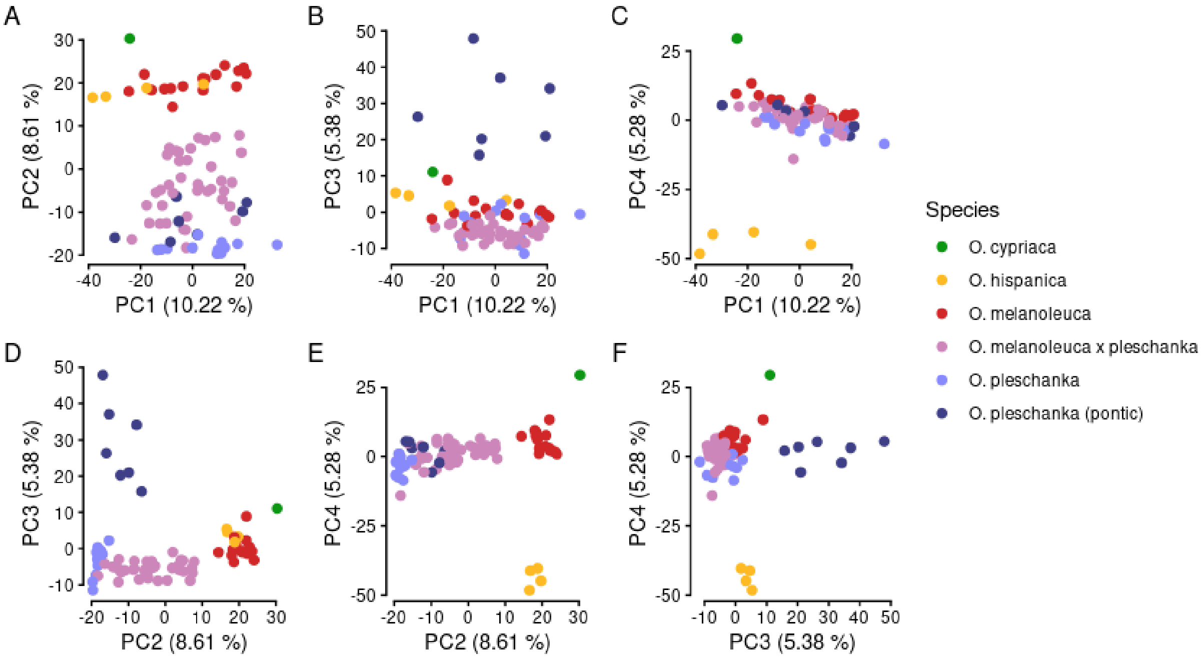


**Figure S2 | PC1 – PC4 of principal component analysis of 436,762 CpG sites.** Percentages in axes labels indicate variance explained by the respective principal component.

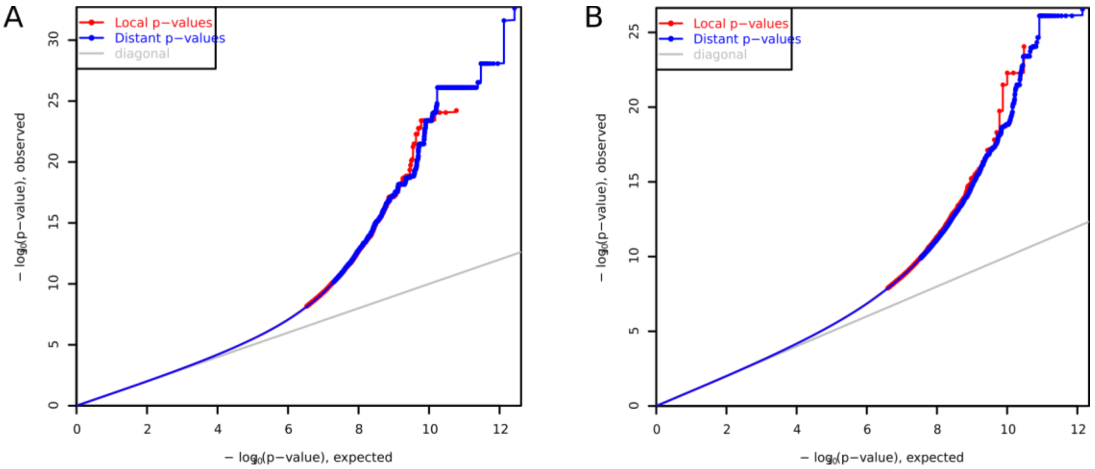


**Figure S3 | QQ-Plot for meQTL analyses for SNPs A) before and B) after LD-pruning.** Plots show the observed vs. expected p-value distribution as returned by matrixEQTL.

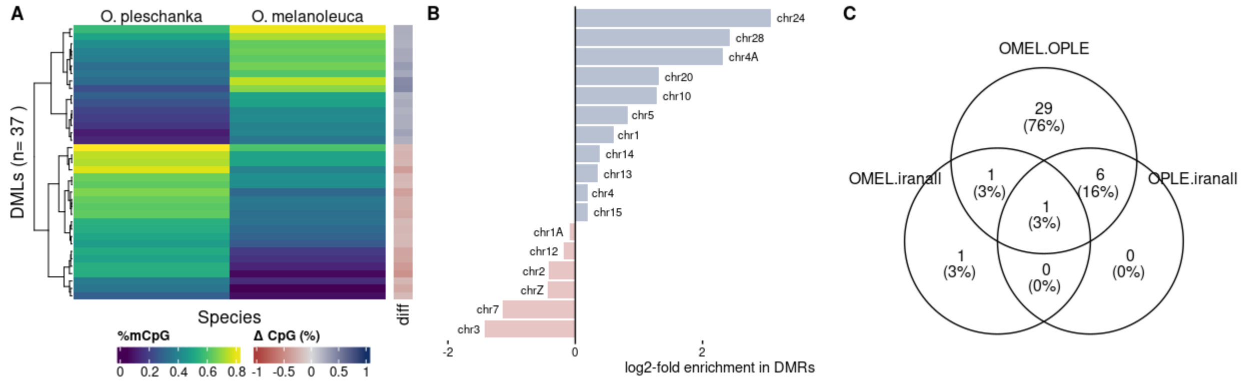


**Figure S4 | Differentially methylated region analysis (DMR). A)** Heatmap of 37 differentially methylated regions called between *O. pleschanka* and *O. melanoleuca*. **B)** Venn diagram of differentially methylated regions called between *O. pleschanka*, *O. melanoleuca*, and hybrids.

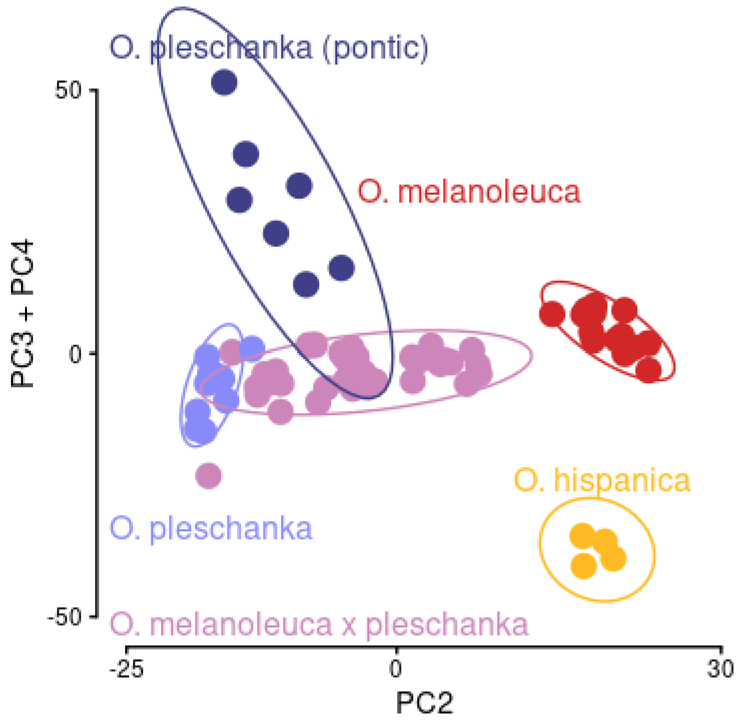


**Figure S5 | PCA with chromosomes 4A and Z removed.** Virtually no difference compared to the PCA using all chromosomes is visible.

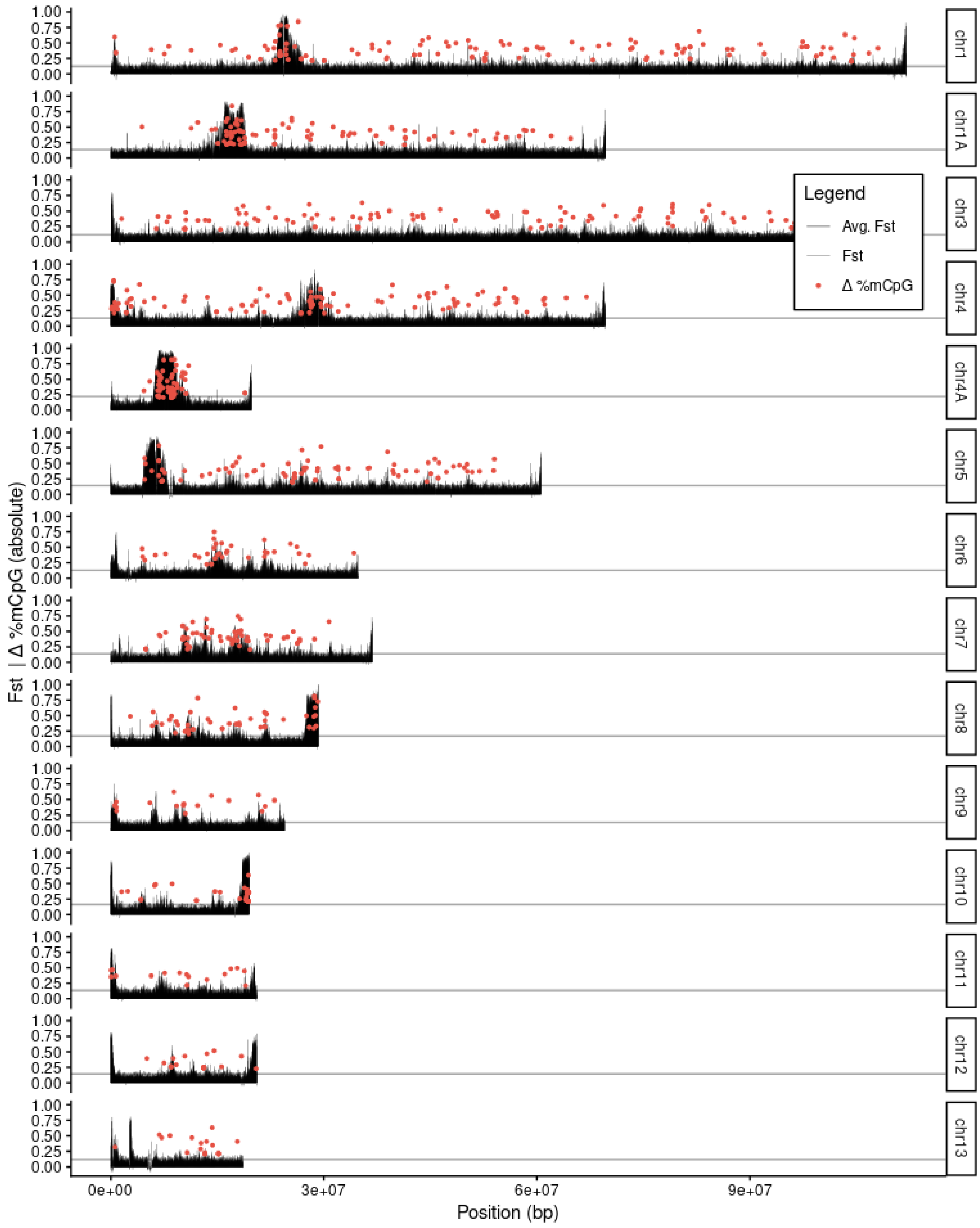


**Figure S6A** | Fixation index ($F_{ST}$) traces and absolute methylation differences between *O. melanoleuca* and *O. pleschanka* for chromosomes 1-13.

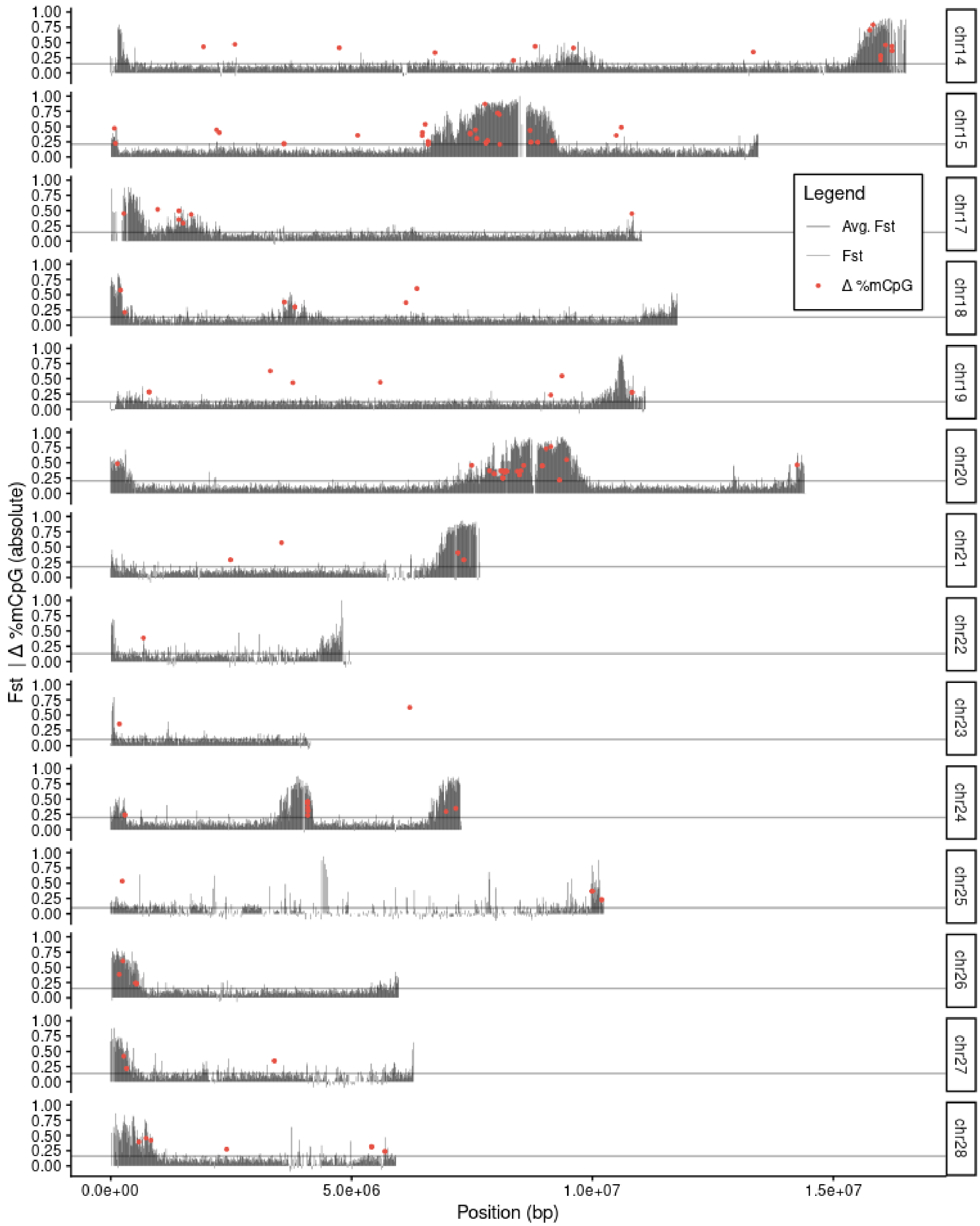


**Figure S6B** | Fixation index ($F_{ST}$) traces and absolute methylation differences between *O. melanoleuca* and *O. pleschanka* for chromosomes 14-28. Note that the assembly for *O. melanoleuca* does not contain chromosome 16. This repeat-rich avian microchromosome can usually not be assembled.

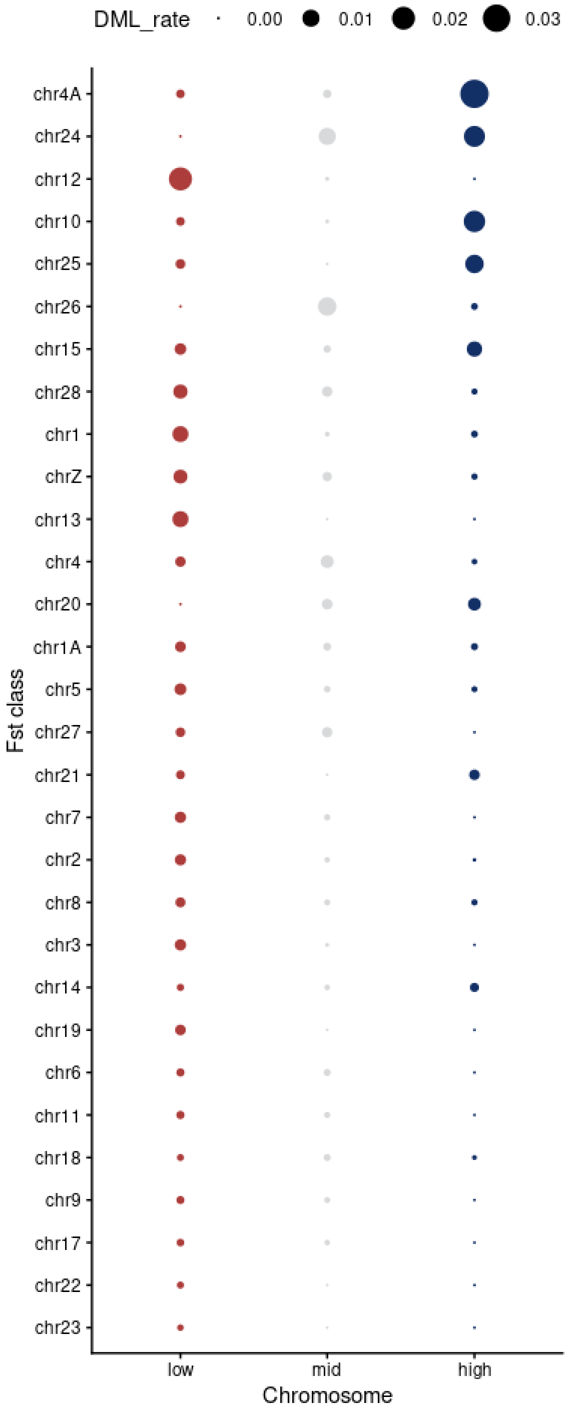


**Figure S7 | DML rate per Fst class per chromosme.** DML rate is calculated as number of $F_{ST}$ windows with DMLs divided by the number of tested CpG sites per chromosome.

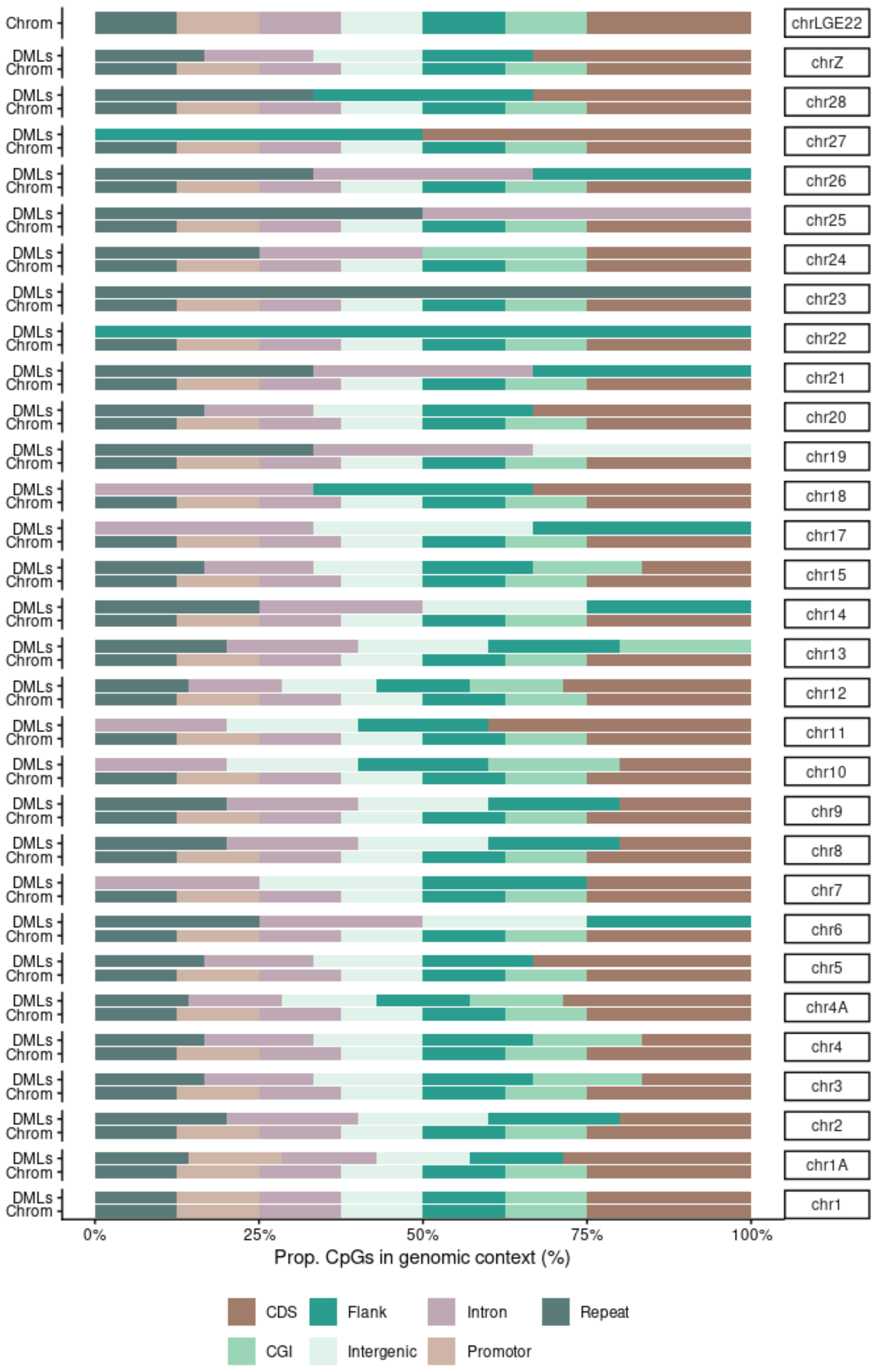


**Figure S8 | Proportional occurence of DMLs per genomic context (DMLs) and chromosomal occurrence of CpG sites per genomic context (Chrom) for each chromosome.**

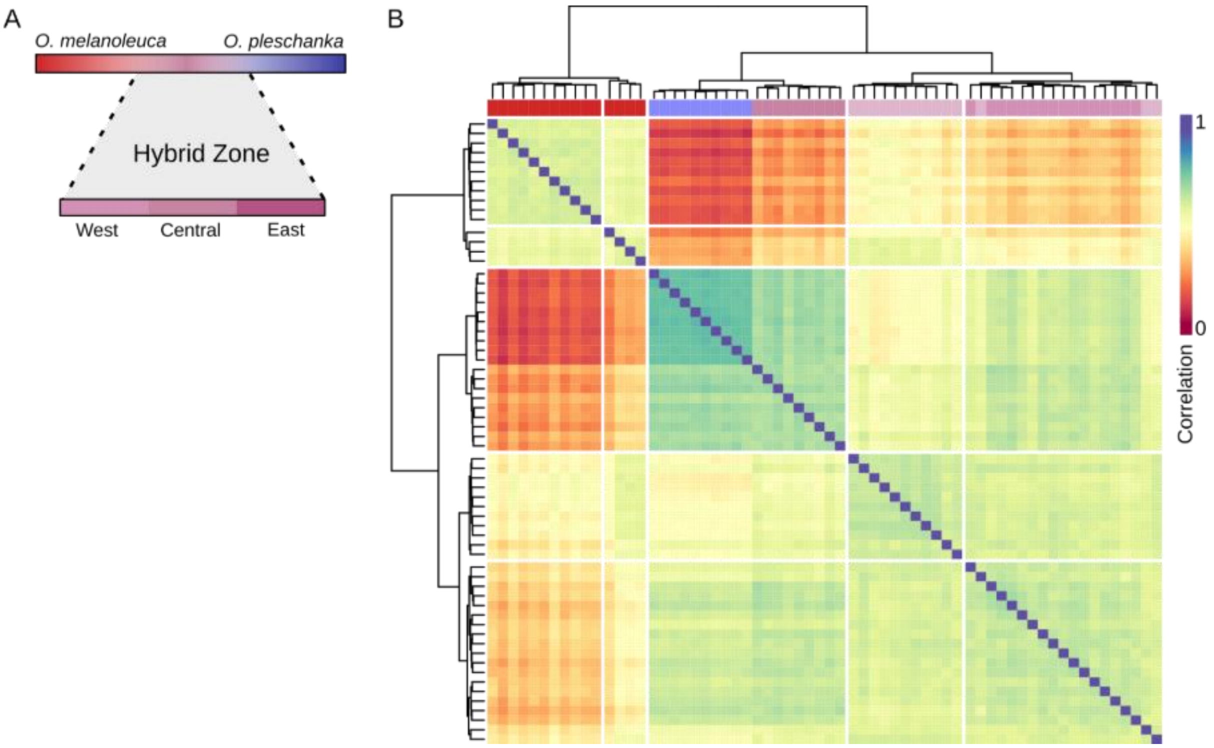


**Figure S9 | Hybrid methylomes cluster into three geographic hybrid-subpopulations. A)** Iranian hybrids were genetically classified into three geographic groups occurring in western, central and eastern Iran. **B)** Unsupervised hierarchical clustering of Spearman correlation for differentiated CpG sites reflect grouping into parental populations and the three hybrid subpopulations with the eastern subpopulations clustering together with parental individuals from *O. pleschanka*.

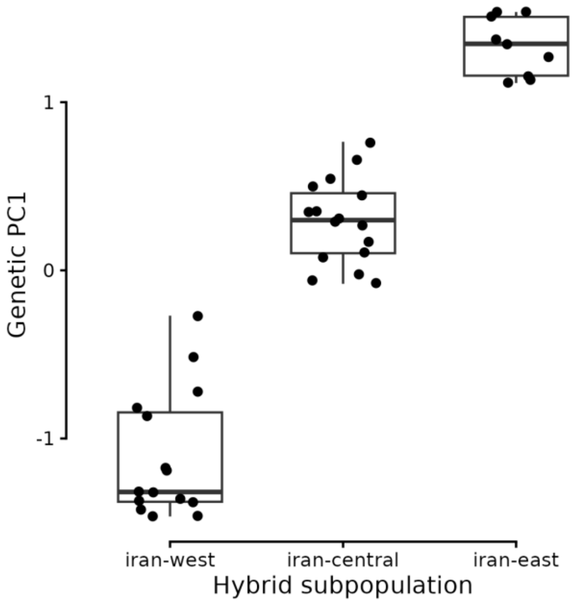


**Figure S10 | Distribution of the first genetically determined principal component (PC1) in Iranian Hybrids showing that central Iranian samples show intermediate ancestry between *O. melanoleuca* and *O. pleschanka*.** Genetically determined PC1 separated *O. melanoleuca* and *O. pleschanka* and forms a gradient for the Iranian hybrid samples from west to east.

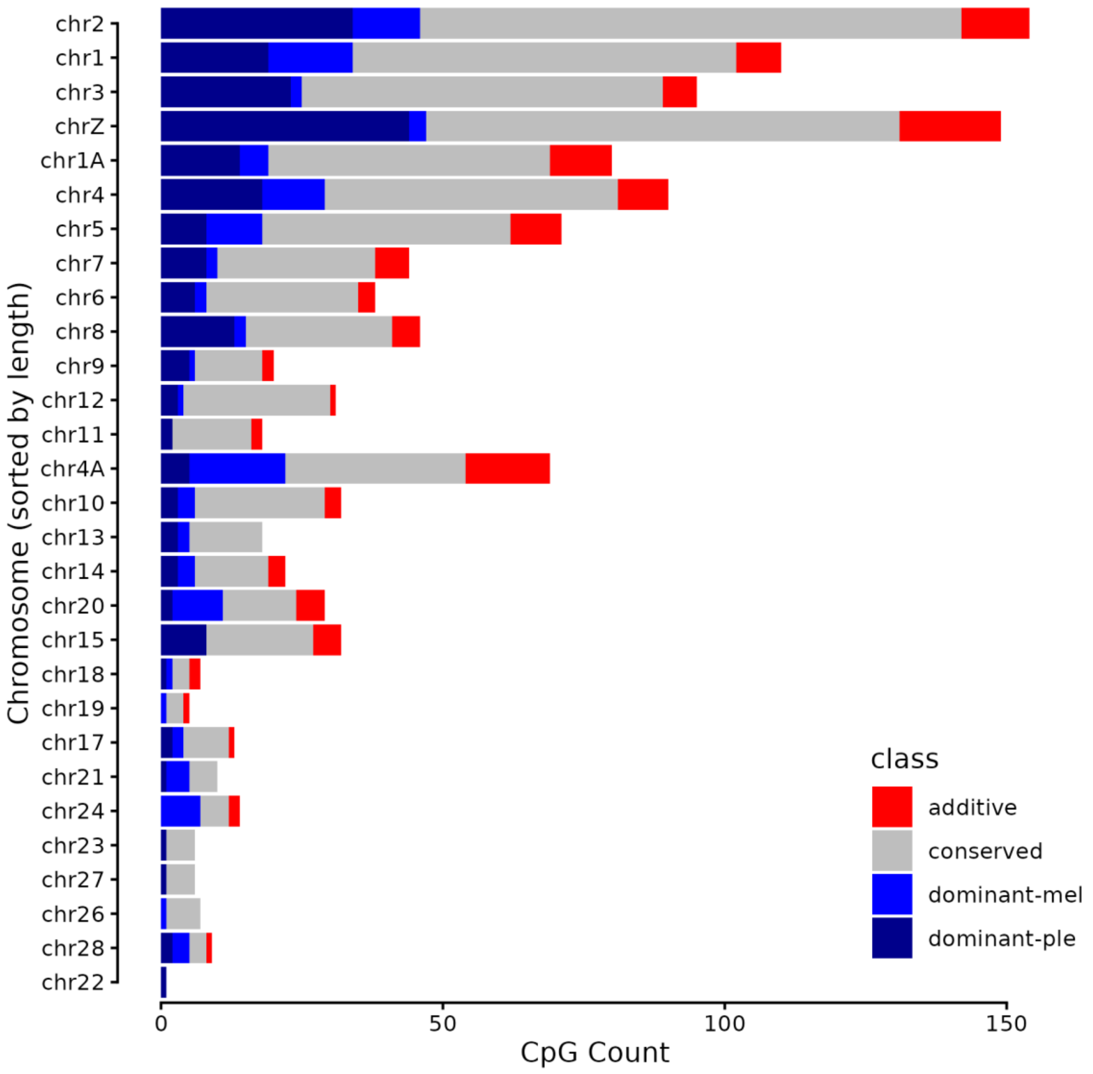


**Figure S11 | Occurrence of classified methylation signatures in Iranian hybrids separated by chromosome.**
